# Deducing the N- and O- glycosylation profile of the spike protein of novel coronavirus SARS-CoV-2

**DOI:** 10.1101/2020.04.01.020966

**Authors:** Asif Shajahan, Nitin T. Supekar, Anne S. Gleinich, Parastoo Azadi

## Abstract

The current emergence of the novel coronavirus pandemic caused by SARS-CoV-2 demands the development of new therapeutic strategies to prevent rapid progress of mortalities. The coronavirus spike (S) protein, which facilitates viral attachment, entry and membrane fusion is heavily glycosylated and plays a critical role in the elicitation of the host immune response. The spike protein is comprised of two protein subunits (S1 and S2), which together possess 22 potential N-glycosylation sites. Herein, we report the glycosylation mapping on spike protein subunits S1 and S2 expressed on human cells through high resolution mass spectrometry. We have characterized the quantitative N-glycosylation profile on spike protein and interestingly, observed unexpected O-glycosylation modifications on the receptor binding domain (RBD) of spike protein subunit S1. Even though O-glycosylation has been predicted on the spike protein of SARS-CoV-2, this is the first report of experimental data for both the site of O-glycosylation and identity of the O-glycans attached on the subunit S1. Our data on the N- and O-glycosylation is strengthened by extensive manual interpretation of each glycopeptide spectra in addition to using bioinformatics tools to confirm the complexity of glycosylation in the spike protein. The elucidation of the glycan repertoire on the spike protein provides insights into the viral binding studies and more importantly, propels research towards the development of a suitable vaccine candidate.

## Introduction

The current major health crisis is caused by the novel severe acute respiratory syndrome coronavirus 2 (SARS-CoV-2) that rapidly spread globally within weeks in early 2020. This highly transmissible infectious disease causes a respiratory illness named COVID-19 (Huang, C., Wang, Y., et al. 2020, Wu, F., Zhao, S., et al. 2020). As of the 31st of March 2020, 750890 cases of COVID-19 and 36405 COVID-19-related deaths have been confirmed globally by the World Health Organization (WHO) (World Health Organization 2020b).

To date, no specific medical treatments or vaccines for COVID-19 have been approved (Li, G.D. and De Clercq, E. 2020, World Health Organization 2020a). Therefore, the scientific community is expending great effort in compiling data regarding the virus, as well as the respiratory illness caused by it, to find effective ways of dealing with this health crisis.

The pathogenic SARS-CoV-2 enters human target cells via its viral transmembrane spike (S) glycoprotein. The spike protein is a trimeric class I fusion protein and consists of two subunits, namely S1 and S2. The S1 subunit facilitates the attachment of the virus, and subsequently the S2 subunit allows for the fusion of the viral and human cellular membranes (Hoffmann, M., Kleine-Weber, H., et al. 2020, Walls, A.C., Park, Y.J., et al. 2020, Zhou, P., Yang, X.L., et al. 2020). The entry receptor for SARS-CoV-2 has been identified as the human angiotensin-converting enzyme 2 (hACE2), and recent studies determined a high binding affinity to hACE2 (Hoffmann, M., Kleine-Weber, H., et al. 2020, Shang, J., Ye, G., et al. 2020, Walls, A.C., Park, Y.J., et al. 2020). Given its literal key role, the S protein is one of the major targets for the development of specific medical treatments or vaccines: neutralizing antibodies targeting the spike proteins of SARS-CoV-2 could prevent the virus from binding to the hACE2 entry receptor and therefore from entering the host cell (Shang, J., Ye, G., et al. 2020).

Each monomer of the S protein is highly glycosylated with 22 predicted N-linked glycosylation sites. Furthermore, four O-glycosylation sites were also predicted (Andersen, K.G., Rambaut, A., et al. 2020). Cryo-electron microscopy (Cryo-EM) provides evidence for the existence of 14–16 N-glycans on 22 potential sites in the SARS-CoV-2 S protein (Walls, A.C., Park, Y.J., et al. 2020). The glycosylation pattern of the spike protein is a crucial characteristic to be considered regarding steric hindrance, chemical properties and even as a potential target for mutation in the future. The N-glycans on S protein play important roles in proper protein folding and priming by host proteases. Since glycans can shield the amino acid residues and other epitopes from cells and antibody recognition, glycosylation can enable the coronavirus to evade both the innate and adaptive immune responses (Walls, A.C., Park, Y.J., et al. 2020, Walls, A.C., Xiong, X., et al. 2019). Elucidating the glycosylation of the viral S protein can aid in understanding viral binding with receptors, fusion, entry, replication and also in designing suitable antigens for vaccine development (Chakraborti, S., Prabakaran, P., et al. 2005, Watanabe, Y., Bowden, T.A., et al. 2019, Zheng, J., Yamada, Y., et al. 2018). Strategies for vaccine development aim to elicit such adaptive immunity through an antibody response at the sites of viral entry (Afrough, B., Dowall, S., et al. 2019).

Here, we report the site-specific quantitative N-linked and O-linked glycan profiling on SARS-CoV-2 subunit S1 and S2 protein through glycoproteomics using high resolution LC-MS/MS. We used recombinant SARS-CoV-2 subunit S1 and S2 expressed in human cells, HEK 293, and observed partial N-glycan occupancy on 17 out of 22 N-glycosylation sites. We found that the remaining five N-glycosylation sites were unoccupied. Remarkably, we have unambiguously identified 2 unexpected O-glycosylation sites at the receptor binding domain (RBD) of subunit S1. O-glycosylation on the spike protein of SARS-CoV-2 is predicted in several recent reports and most of these predictions are for sites in proximity to furin cleavage site (S1/S2) as similar sites are O-glycosylated in SARS-CoV-1 (Andersen, K.G., Rambaut, A., et al. 2020). However, we observed O-glycosylation at two sites on the RBD of spike protein subunit S1, and this is the first report on the evidence for such glycan modification at a crucial viral attachment location. Site-specific analysis of N- and O-glycosylation information of SARS-CoV-2 spike protein provides basic understanding of the viral structure, crucial for the identification of immunogens for vaccine design. This in turn has the potential of leading to future therapeutic intervention or prevention of COVID-19.

## Results

Studies over the past two decades have shown that glycosylation on the protein antigens can play crucial roles in the adaptive immune response. Thus, it is obvious that the glycosylation on the protein antigen is relevant for the development of vaccines, and it is widely accepted that the lack of information about the glycosylation sites hampers the design of such vaccines (Wolfert, M.A. and Boons, G.J. 2013).

### Mapping N-glycosylation on SARS-CoV-2 spike protein

We have procured culture supernatants of HEK 293 cells expressing SARS-CoV-2 subunit 1 and subunit 2 separately. The proteins were expressed with a His tag with Val16 to Gln690 for subunit 1 and Met697 to Pro1213 for subunit 2. According to manufacturers, SDS-PAGE of the proteins showed a higher molecular weight than the predicted 75 and 60 kDa, respectively, due to glycosylation. Since the proteins were unpurified, we fractionated them through SDS-PAGE on separate lanes and cut the bands corresponding to subunit 1 and subunit 2. The gels were stained with Coomassie dye, and gel bands were cut into small pieces, de-stained, reduced, alkylated and subjected to in-gel protease digestion. We employed trypsin, chymotrypsin, and both trypsin-chymotrypsin in combination to generate glycopeptides that contain a single N-linked glycan site. The glycopeptides were further analyzed by high resolution LC-MS/MS, using a glycan oxonium ion product dependent HCD triggered CID program. The LC-MS/MS data were analyzed using Byonic software, each detected spectrum was manually validated and false detections eliminated.

We identified the glycan compositions at 17 out of the 22 predicted N-glycosylation sites of the SARS-CoV-2 S1 and S2 proteins and found the remaining five sites unoccupied (Figure 2, 3). We observed both high mannose and complex-type glycans across the N-glycosylation sites but found no hybrid type N-glycans. We quantified the relative intensities of glycans at each site by comparing the area under the curve of each glycopeptide peak on the LC-MS chromatogram. A recent preprint investigated the N-glycosylation on SARS-CoV-2 and reported prevalence of hybrid-type glycans (Watanabe, Y., Allen, J.D., et al. 2020). In contrast, we observed a combination of high mannose and complex-type, but no hybrid-type glycans on most of the sites. We discovered predominantly highly processed sialylated complex-type glycans on sites N165, N282, N801, and N1098 (Figure 3, 4). The highly sialylated glycans at N234 and N282, which are at the RBD can act as determinant in viral binding with hACE2 receptors (Hoffmann, M., Kleine-Weber, H., et al. 2020, Tortorici, M.A., Walls, A.C., et al. 2019, Walls, A.C., Park, Y.J., et al. 2020). Similar to one recent report, we observed Man5GlcNAc2 as a predominant structure across all sites (Watanabe, Y., Allen, J.D., et al. 2020). However, we observed significantly unoccupied peptides on both S1 and S2 subunits. Sites N17, N603, N1134, N1158 and N1173 were completely unoccupied, although further studies with higher concentration and purity of proteins are required to validate this finding (Figure 1, 2). On subunit S2, the detection of N-glycosylation at sites N709, N717, and N1134 was ambiguous as the quality of the MS/MS spectra was not satisfactory and we are currently evaluating the possibilities of other post translational modifications adjacent to these sites.

**Figure 1:**
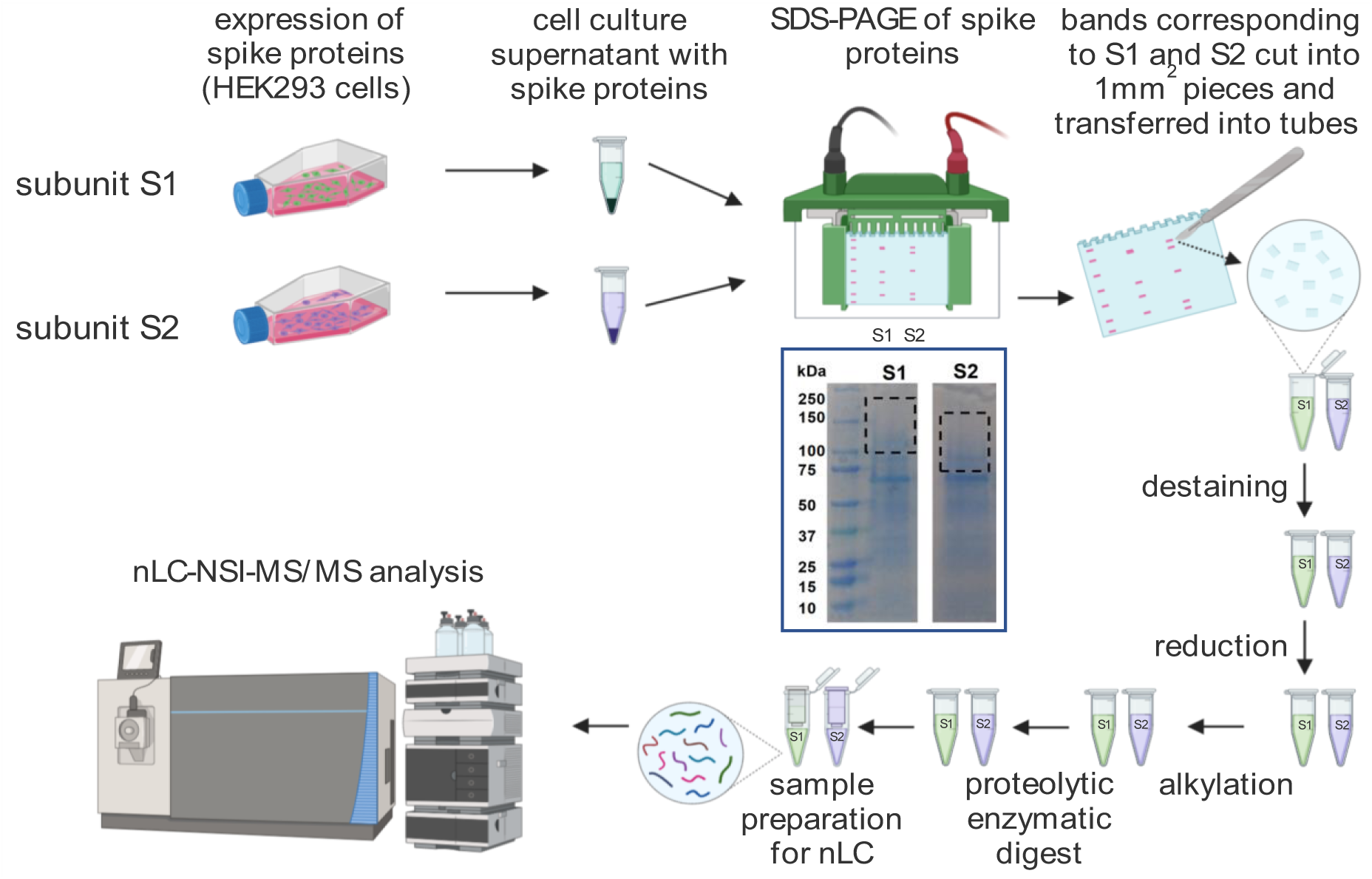
The SARS-Cov-2 spike proteins recombinatly expressed on HEK293 supernatant were fractionated through SDA-PAGE, subsequently digested by proteases anc analyzed by nLC-NSI-MS/MS. The expression of SARS-Cov-2 spike protein subunits S1 and S2 on HEK 293 culture supernatant showed higher molecular weight upon SDS-PAGE than expected, because of glycosylation. Thus, the gel bands corresponding to the molecular weight of 200 kDa to 100 kDa for S1 and 150 kDa to 80 kDa for S2 were cut, proteins were lysed after reduction-alkylation and analyzed by LC-MS/MS (created with biorender.com).

**Figure 2:**
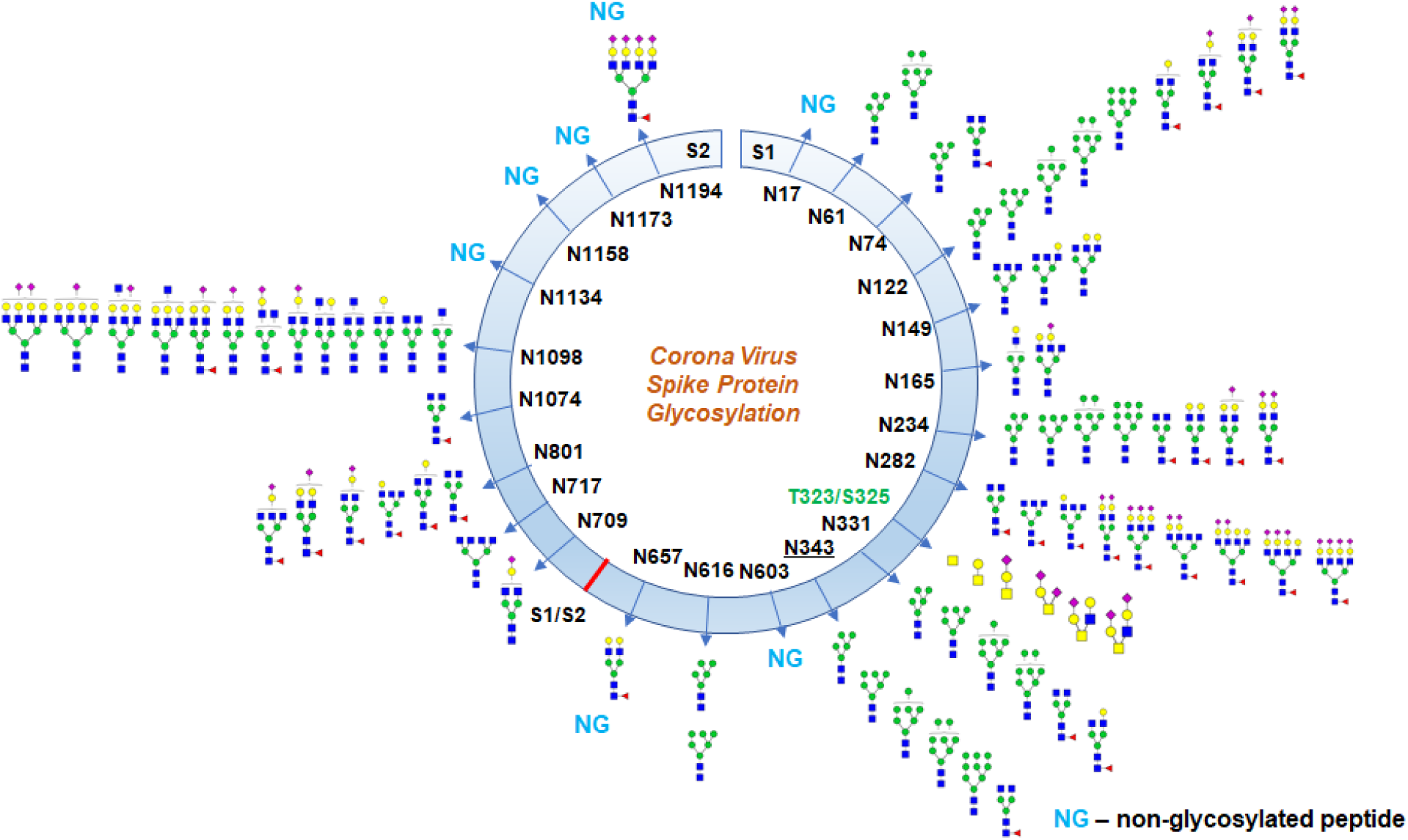
Glycosylation profile on coronavirus SARS-CoV-2 characterized by high-resolution LC-MS/MS. About 17 N-glycosylation sites were found occupied out of 22 potential sites along with two O-glycosylation sites bearing core-1 type O-glycans. Some N-glycosylation sites were partially glycosylated.

**Figure 3:**
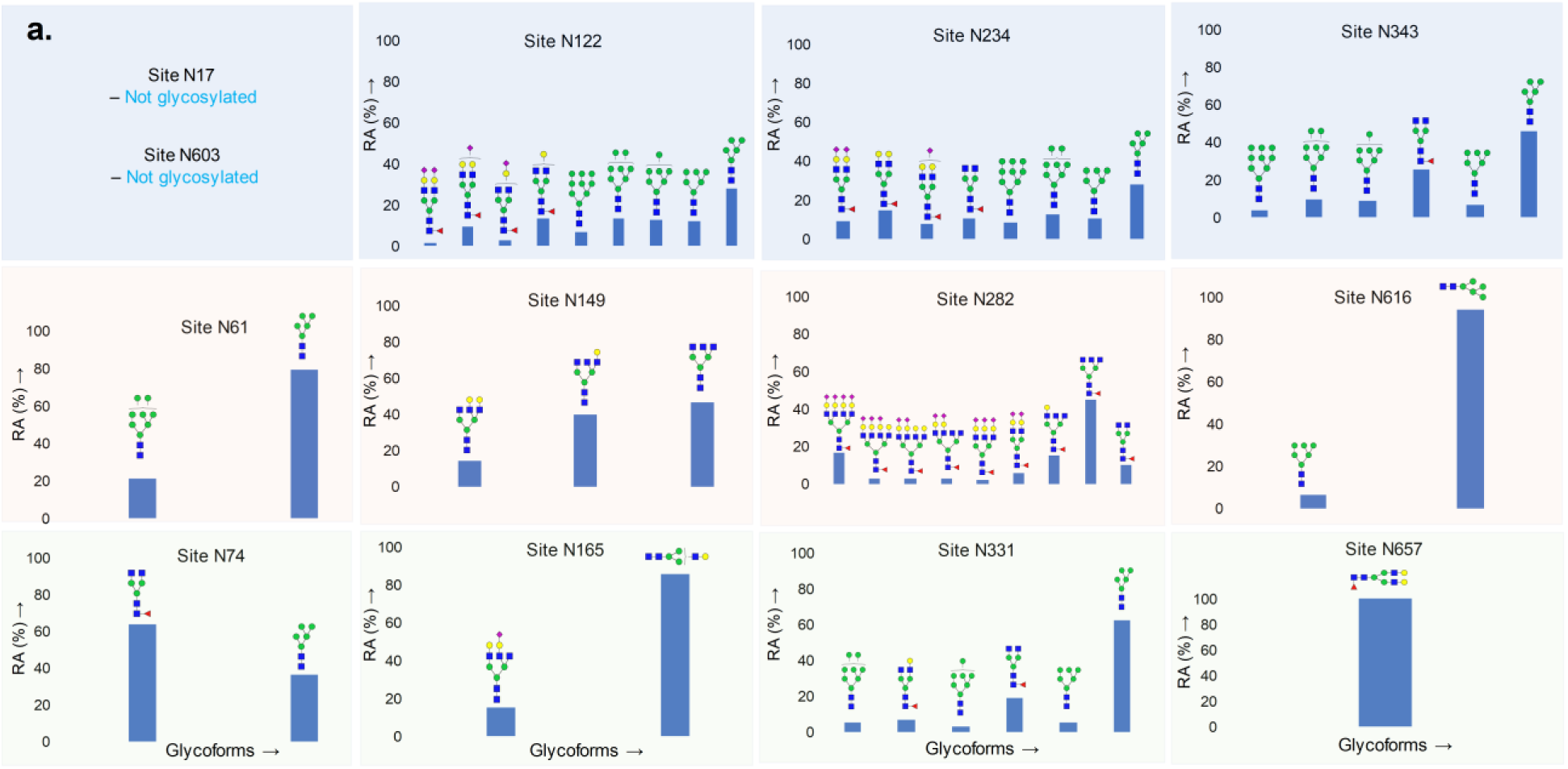
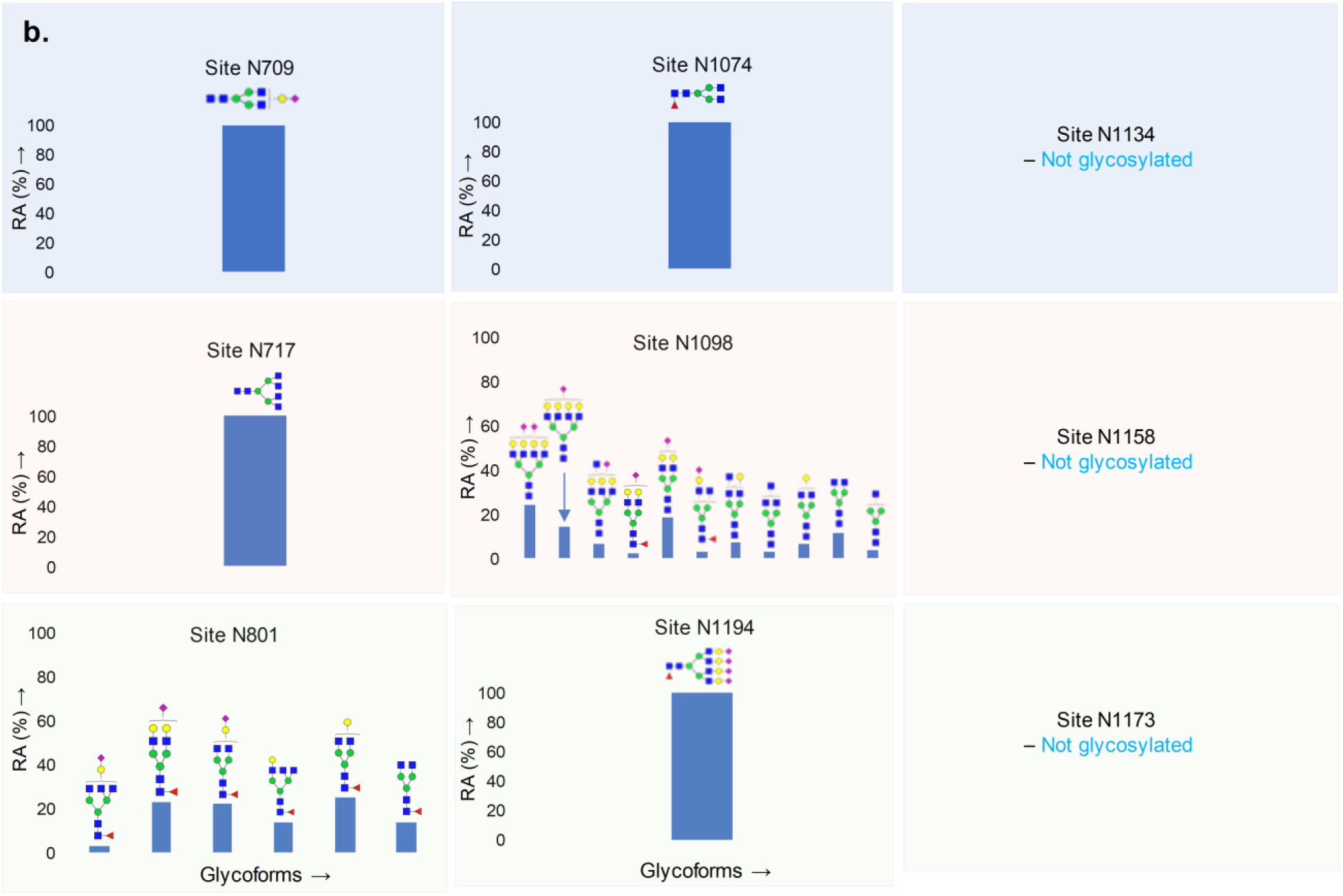
Quantitative glycosylation profile of N-glycans on coronavirus SARS-CoV-2 spike protein characterized by high-resolution LC-MS/MS. **a.** 13 sites on subunit S1; **b.** 9 sites on subunit S2. RA – Relative abundances.

**Figure 4:**
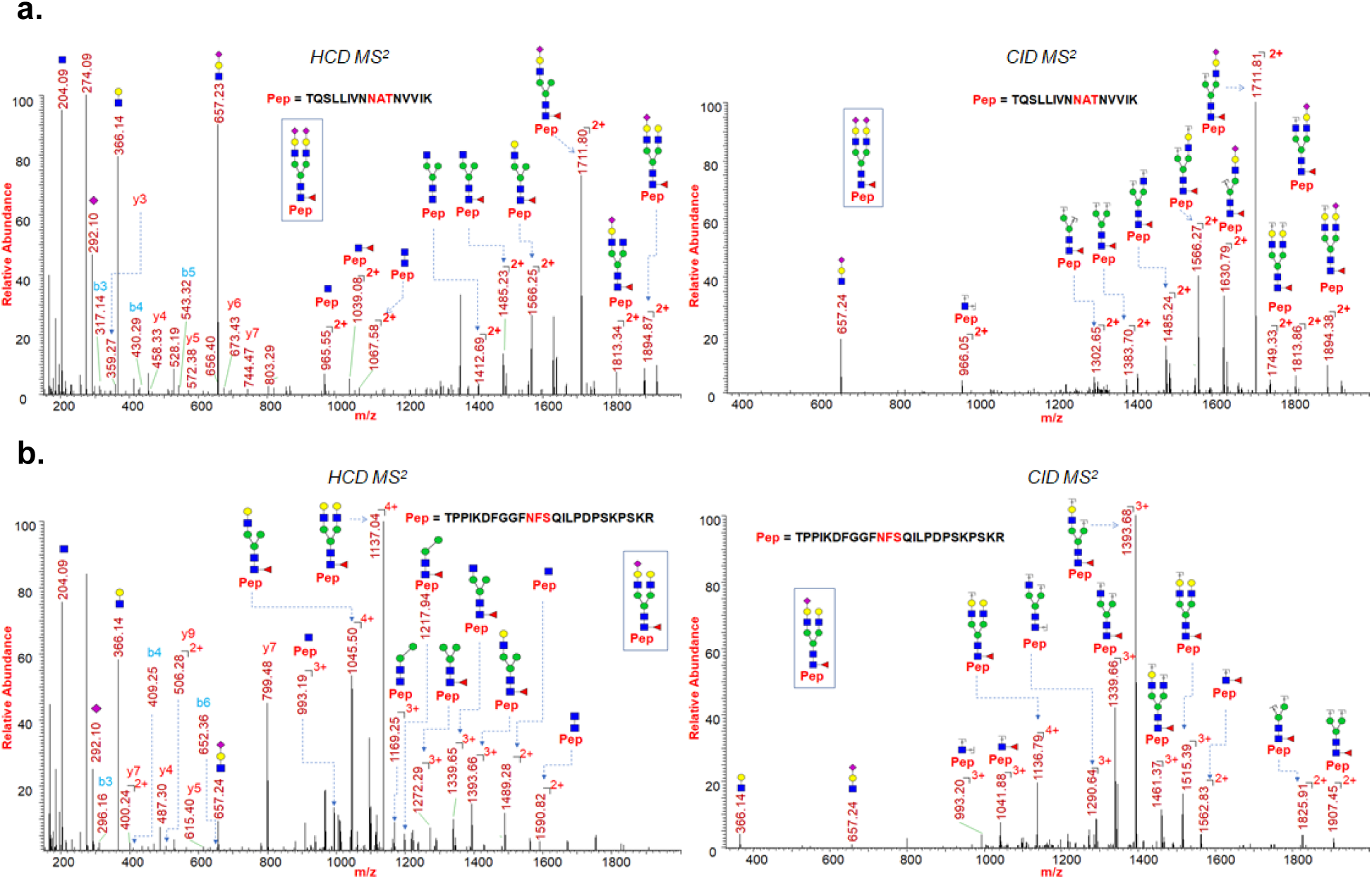
HCD and CID MS/MS spectra showing glycan neutral losses, oxonium ions and peptide fragments of **a.** representative N-glycopeptide TQSLLIVNNATNVVIK (site N122) of spike protein subunit 1; **b.** representative N-glycopeptide TPPIKDFGGFNFSQILPDPSKPSKR (site N801) of spike protein subunit 2.

### Identification of O-Glycosylation on SARS-CoV-2 spike protein

We have evalulated O-glycosylation on the S1 and S2 subunits of SARS-CoV-2 spike protein by searching LC-MS/MS data for common O-glycosylation modifications. Interestingly, our O-glycoproteomic profiling indicated O-glycosylation at sites Thr323 and Ser325 on the S1 subunit of SARS-CoV-2 spike protein (Figure 5, 6). Since O-glycosylation at Thr323 and Ser325 on the spike protein has not been reported before and is not indicated based on cryo-electron microscopy data of SARS-CoV-2 S protein, we evaluated the detected O-glycopeptide manually (Walls, A.C., Park, Y.J., et al. 2020). We observed very strong evidence for the presence of O-glycosylation at site Thr323 as b and y ions of the peptide 320VQPTESIVR328 with high mass accuracy. Upon manual validation of the fragment ions of O-glycopeptide 320VQPTESIVR328 we observed that Thr323 is the predominantly occupied site. This conclusion was based on the b (b1-m/z 228.13) and y (y1-m/z 175.12, y2-m/z 274.19, y4-474.30, y5-603.34 m/z, and y7+glycan-1748.77 m/z) ions we detected upon fragmentation of the glycopeptide (Figure 6a). In addition, neutral losses and the detection of oxonium ions also confirmed the presence of glycosylation on these peptides. Core-1 mucin type O-glycans such as GalNAc, GalNAcGal and GalNAcGalNeuAc2 and Core-2 glycans GalNAcGalNeuAc(GlcNAcGal) and GalNAcGalNeuAc(GlcNAcGalNeuAc) were observed on site Thr323 (Figure 6). Possibly, Ser325 is occupied with HexNAcHexNeuAc glycan together with T323 (Figure 5), but this could not be confirmed unambiguously as Electron Transfer Dissociation (ETD) fragmentation on this peptide was not successful because of lower charge states of peptide (Supp Fig S3, S4) (Shajahan, A., Heiss, C., et al. 2017).

**Figure 5:**
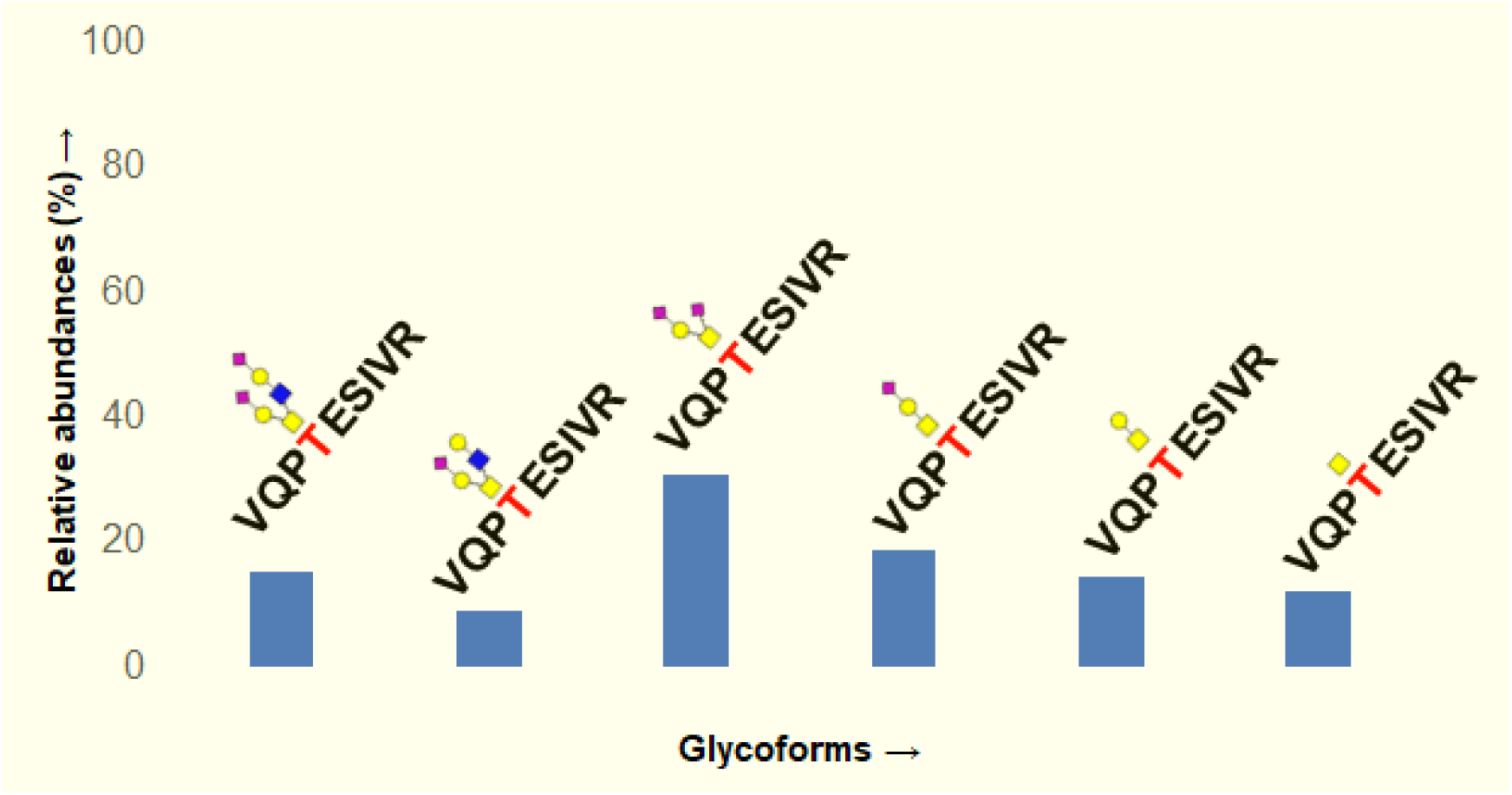
Quantitative glycosylation profile of O-glycans on sites T323 of coronavirus SARS-Cov-2 spike protein characterized by high-resolution LC-MS/MS.

**Figure 6:**
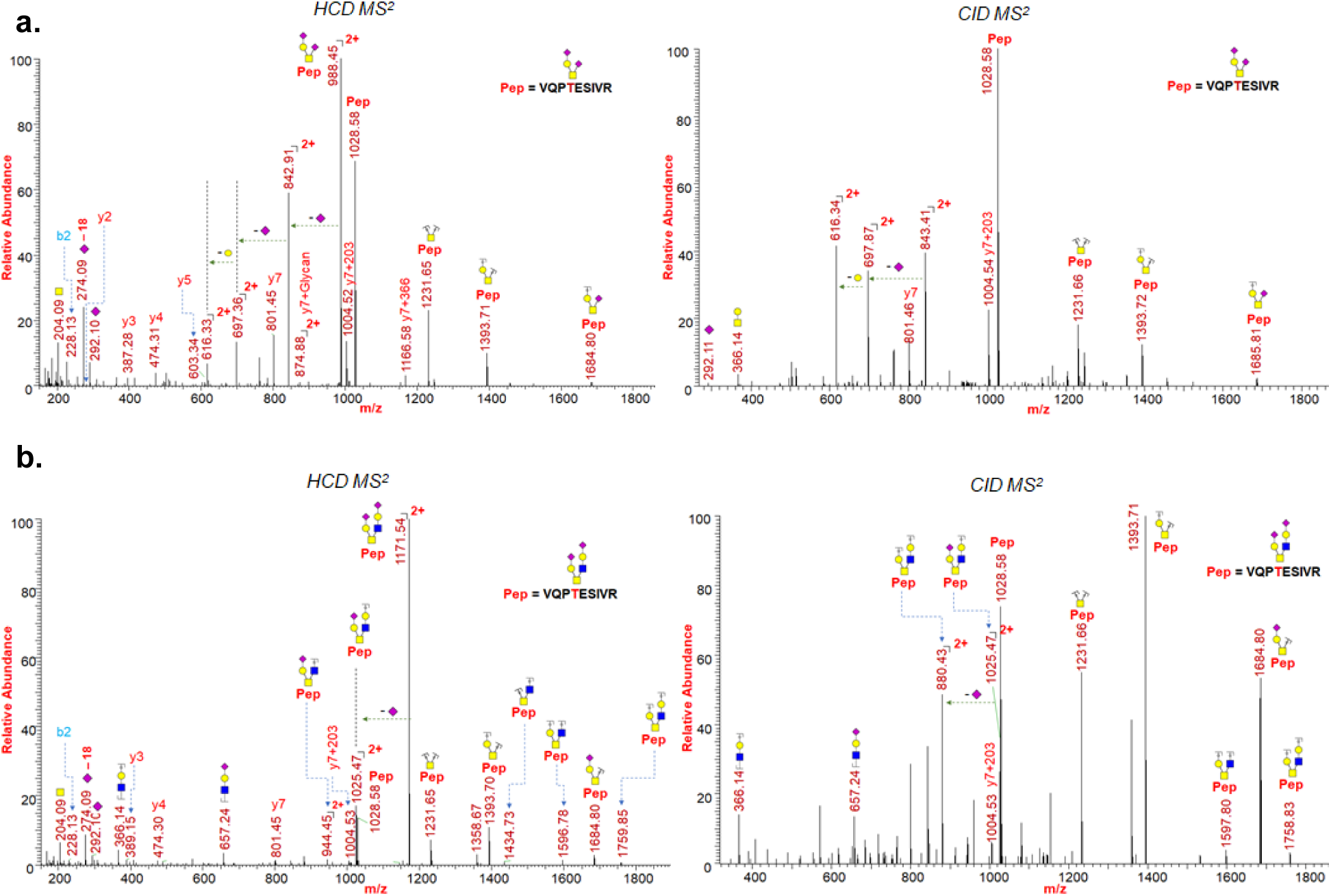
HCD and CID MS/MS spectra showing glycan neutral losses, oxonium ions and peptide fragments of **a.** representative O-Glycopeptide ^320^VQPTESIVR^328^ with core 1 type GalNAcGalNeuAc2 glycan detected on site Thr323 of spike protein subunit 1; **b.** representative O-Glycopeptide ^320^VQPTESIVR^328^ with core 2 type GalNAcGalNeuAc(GlcNAcGalNeuAc) glycan detected on site Thr323 of spike protein subunit 1.

## Discussion

Two very recent preprints reporting N-glycosylation on spike protein of SARS-CoV-2 showed different glycosylation profiles, and both reports were different from our results although all three studies utilized recombinant S protein from an HEK 293-based expression system (Watanabe, Y., Allen, J.D., et al. 2020, Zhang, Y., Zhao, W., et al. 2020). This indicates the caution to be exercised in determining the glycosylation on viral antigens generated from various sources as the changes in glycosylation pattern can influence the efficacy of potential vaccine candidates (Chang, D. and Zaia, J. 2019).

O-glycans, which are involved in protein stability and function, have been observed on some viral proteins and have been suggested to play roles in the biological activity of viral proteins (Andersen, K.G., Rambaut, A., et al. 2020, Bagdonaite, I. and Wandall, H.H. 2018). A comparative study of human SARS-CoV-2 S protein with other coronavirus S proteins has shown that Ser673, Thr678, and Ser686 are conserved O-glycosylation locations, and SARS-CoV-2 S1 protein was suggested to be O-glycosylated at these locations (Andersen, K.G., Rambaut, A., et al. 2020). Although it is unclear what function these predicted O-linked glycans perform, they have been suggested to create a ‘mucin-like domain’ which could shield SARS-CoV-2 spike protein epitopes or key residues (Bagdonaite, I. and Wandall, H.H. 2018). Since some viruses can utilize mucin-like domains as glycan shields for immunoevasion, researchers have highlighted the importance of experimental studies for the determination of predicted O-linked glycosylation sites (Andersen, K.G., Rambaut, A., et al. 2020, Bagdonaite, I. and Wandall, H.H. 2018). We evaluated the O-glycosylation site prediction using the widely accepted tool Net-O-Gly server 4.0 (Steentoft, C., Vakhrushev, S.Y., et al. 2013). However, the tool did not find any strong prediction for the O-glycosylation except for the sites Ser673, Thr678 and Ser686.

Our analysis confirmed the presence of O-glycosylation at Thr323 and indicated possible glycosylation at Ser325 (Figure 5, 6). Intriguingly, the possible O-glycosylation at Thr323 of SARS-CoV-2 subunit 1 glycoprotein has been predicted by computational analysis in a very recent preprint report (Uslupehlivan, M. and Sener, E. 2020). The accuracy of our observation of O-glycosylation at Thr323 is further confirmed by the presence of proline at location 322, considering the well-established fact that the frequency of occurrence of proline residues is higher adjacent to O-glycosylation sites (Thanka Christlet, T.H. and Veluraja, K. 2001). Cryo-EM studies on SARS-CoV-2 indicate that the binding of S protein to the hACE2 receptor primarily involves extensive polar residue interactions between RBD and the peptidase domain of hACE2 (Hoffmann, M., Kleine-Weber, H., et al. 2020, Walls, A.C., Park, Y.J., et al. 2020). The S protein RBD located in the S1 subunit of SARS-CoV-2 undergoes a hinge-like dynamic movement to enhance the capture of the receptor RBD with hACE2, displaying 10–20-fold higher affinity for the human ACE2 receptor than SARS-CoV-1, which partially explains the higher transmissibility of the new virus (Wrapp, D., Wang, N., et al. 2020, Yan, R., Zhang, Y., et al. 2020). The residues Thr323 and Ser325 are located at the RBD of the S1 subunit of SARS-Cov-2, and thus the O-glycosylation at this location could play a critical role in viral binding with hACE2 receptors (Andersen, K.G., Rambaut, A., et al. 2020, Hoffmann, M., Kleine-Weber, H., et al. 2020). Our observation will pave the way for future studies to understand the implication of O-glycosylation at the RBD of S1 protein in viral attachment with hACE2 receptors.

Our comprehensive N- and O-glycosylation characterization of SARS-CoV-2 expressed in a human cell system provides insights into site-specific N- and O-glycan decoration on the trimeric spike protein. We have employed an extensive manual interpretation strategy for the assignment of each glycopeptide structure in order to eliminate possibilities of ambiguous software based annotation. We are currently working on elucidating other potential post-translational modifications on SARS-CoV-2 spike protein as understanding the protein modifications in detail is important to guide future researches on disease interventions involving spike protein. Detailed glycan analysis is important for the development of glycoprotein-based vaccine candidates as a means to correlate the structural variation with immunogenicity. Glycosylation can serve as a measure to evaluate antigen quality as various expression systems and production processes are employed in vaccine manufacture. The understanding of complex sialylated N-glycans and sialylated mucin type O-glycans, particularly in the RBD domain of the spike protein of SARS-CoV-2, provides basic knowledge useful for elucidating the viral infection pathology in future therapeutic possibilities, as well as in the design of suitable immunogens for vaccine development.

## Materials and methods

Dithiothreitol (DTT) and iodoacetamide (IAA) were purchased from Sigma Aldrich (St. Louis, MO). Sequencing-grade modified trypsin and chymotrypsin were purchased from Promega (Madison, WI). All other reagents were purchased from Sigma Aldrich unless indicated otherwise. Data analysis was performed using Byonic 3.5 software and manually using Xcalibur 4.2. The SARS-CoV-2 spike protein subunit 1 (Cat No. 230-20407) and subunit 2 (Cat No. 230-20408) were purchased from RayBiotech (Atlanta, GA).

### Protease digestion and extraction of peptides from SDS-PAGE

The protein subunits S1 and S2 as HEK 293 culture supernatants were fractionated on separate lanes using SDS-PAGE. The gel was stained by Coomassie dye and the bands corresponding to subunit 1 (200 to 100 kDa) and subunit 2 (150 to 80 kDa) were cut into smaller pieces (1 mm squares approx.) and transferred to clean tubes. The gel pieces were de-stained by adding 100 µL acetonitrile (ACN): 50mM NH_4_HCO_3_ (1:1) and incubated at room temperature (RT) for about 30 min. Tubes were centrifuged, the supernatant was discarded, and 100 µL ACN was added before incubation for 30 min. The proteins on gel pieces were reduced by adding 350 µL 25 mM DTT and incubating at 60 °C for 30 min. The tubes were cooled to RT and the supernatant removed. The gels were washed with 500 µL of ACN, 350 µL 90 mM IAA was added, and the mixture was incubated at RT for 20 min in the dark. Proteins were digested by adding sequence-grade trypsin and/or chymotrypsin (we performed digests with both enzymes individually and as a cocktail) in digestion buffer (50 mM NH_4_HCO_3_) for 18 h at 37 °C separately. The peptides were extracted out from the gel by addition of 1:2 H2O:ACN containing 5% formic acid (500 µL), and the released peptides were speed-dried. The samples were reconstituted in aqueous 0.1% formic acid for LC-MS/MS experiments.

### Data acquisition of protein digest samples using nano-LC-MS/MS

The glycoprotein digests were analyzed on an Orbitrap Fusion Tribrid mass spectrometer equipped with a nanospray ion source and connected to a Dionex binary solvent system (Waltham, MA). Pre-packed nano-LC columns of 15 cm length with 75 µm internal diameter (id), filled with 3 µm C18 material (reverse phase) were used for chromatographic separation of samples. The precursor ion scan was acquired at 120,000 resolution in the Orbitrap analyzer and precursors at a time frame of 3 s were selected for subsequent MS/MS fragmentation in the Orbitrap analyzer at 15,000 resolution. The LC-MS/MS runs of each digest were conducted for both 72 min and 180 min in order to separate the glycopeptides. The threshold for triggering an MS/MS event was set to 1000 counts, and monoisotopic precursor selection was enabled. MS/MS fragmentation was conducted with stepped HCD (Higher-energy Collisional Dissociation) product triggered CID (Collision-Induced Dissociation) (HCDpdCID) program. Charge state screening was enabled, and precursors with unknown charge state or a charge state of +1 were excluded (positive ion mode). Dynamic exclusion was enabled (exclusion duration of 30 secs).

### Data analysis of glycoproteins

The LC-MS/MS spectra of tryptic, chymotryptic and combined tryptic /chymotryptic digests of glycoproteins were searched against the .fasta sequence of spike protein S1 and S2 subunit using the Byonic software by choosing appropriate peptide cleavage sites (semi-specific cleavage option enabled). Oxidation of methionine, deamidation of asparagine and glutamine, possible common human N-glycans and O-glycan masses were used as variable modifications. The LC-MS/MS spectra were also analyzed manually for the glycopeptides with the support of the Xcalibur software. The HCDpdCID MS2 spectra of glycopeptides were evaluated for the glycan neutral loss pattern, oxonium ions and glycopeptide fragmentations to assign the sequence and the presence of glycans in the glycopeptides.

## Acknowledgements

Financial support from the US National Institutes of Health (S10OD018530) is gratefully acknowledged. This work was also supported in part by the U.S. Department of Energy, Office of Science, Basic Energy Sciences, under Award DE-SC0015662 to DOE - Center for Plant and Microbial Complex Carbohydrates at the Complex Carbohydrate Research Center, USA.

## Abbreviations

S: Spike
RBD: receptor binding domain
COVID-19: coronavirus disease
hACE2: human angiotensin-converting enzyme 2
HEK: human embryonic kidney
SDS-PAGE: sodium dodecyl sulfate - polyacrylamide gel electrophoresis
DTT: dithiothreitol
IAA: iodoacetamide
ACN: acetonitrile
HCD: Higher-energy Collisional Dissociation
pd: product triggered
CID: Collision-Induced Dissociation
ETD: Electron Transfer Dissociation

## Contributions

A.S. and P.A. conceived of the paper; A.S., N.S. and A.G. contributed equally and performed experiments; everyone contributed toward writing the paper; P.A. monitored the project.

## Competing interests

The authors certify that they have no competing interests.

## Supplementary material

Annotated MS/MS spectra are incorporated as a separate supplementary material file.

