## Supplementary material for "Deducing the N- and O- glycosylation profile of the spike protein of novel coronavirus SARS-CoV-2"

### Supplementary Information

Table of Contents

| Description | Page No. |
|---|---|
| Supp. Figure S1-S4: Spectra of intact O-glycopeptide with assigned O-glycans at T323 & S325 | S-2 to S-3 |
| Supp. Figure S5: Spectrum of non-glycosylated peptide at N17 | S-4 |
| Supp. Figure S6: Spectrum of intact N-glycopeptide with assigned N-glycan at N61 | S-4 |
| Supp. Figure S7: Spectrum of intact N-glycopeptide with assigned N-glycans at N74 | S-5 |
| Supp. Figure S8: Spectrum of intact N-glycopeptide with assigned N-glycans at N122 | S-5 |
| Supp. Figure S9: Spectrum of intact N-glycopeptide with assigned N-glycan at N149 | S-6 |
| Supp. Figure S10: Spectrum of intact N-glycopeptide with assigned N-glycan at N165 | S-6 |
| Supp. Figure S11: Spectrum of intact N-glycopeptide with assigned N-glycan at N234 | S-7 |
| Supp. Figure S12: Spectrum of intact N-glycopeptide with assigned N-glycan at N282 | S-7 |
| Supp. Figure S13: Spectrum of intact N-glycopeptide with assigned N-glycan at N331 | S-8 |
| Supp. Figure S14: Spectrum of intact N-glycopeptide with assigned N-glycan at N343 | S-8 |
| Supp. Figure S15: Spectrum of non-glycosylated peptide at N603 | S-9 |
| Supp. Figure S16: Spectrum of intact N-glycopeptide with assigned N-glycan at N616 | S-9 |
| Supp. Figure S17: Spectrum of intact N-glycopeptide with assigned N-glycan at N657 | S-10 |
| Supp. Figure S18: Spectrum of intact N-glycopeptide with assigned N-glycan at N801 | S-10 |
| Supp. Figure S19: Spectrum of intact N-glycopeptide with assigned N-glycan at N1074 | S-11 |
| Supp. Figure S20: Spectrum of intact N-glycopeptide with assigned N-glycan at N1098 | S-11 |
| Supp. Figure S21: Spectrum of non-glycosylated peptide at N1158 | S-12 |
| Supp. Figure S22: Spectrum of non-glycosylated peptide at N1173 | S-12 |
| Supp. Figure S23: Spectrum of intact N-glycopeptide with assigned N-glycan at N1194 | S-13 |

Glycopeptide Analysis:

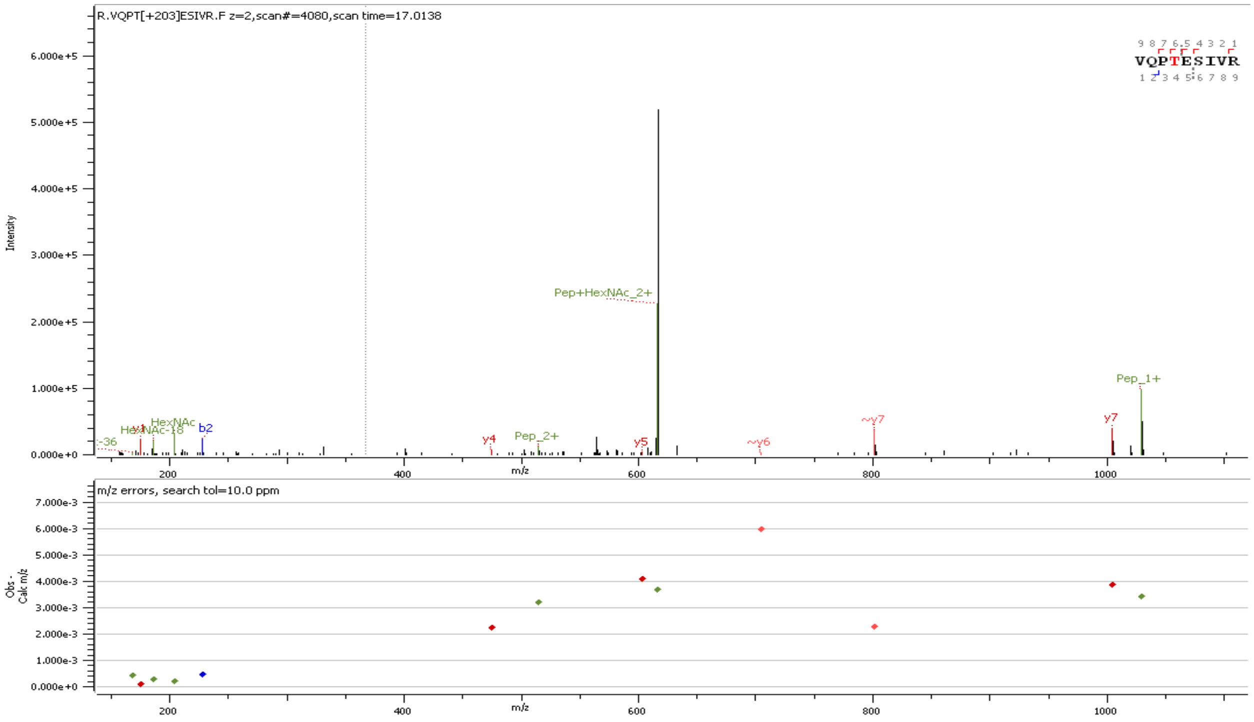


**Supp. Figure S1** Representative HCD MS/MS spectrum of intact O-glycopeptide with assigned O-glycan ($GalNAc_1$) at T323

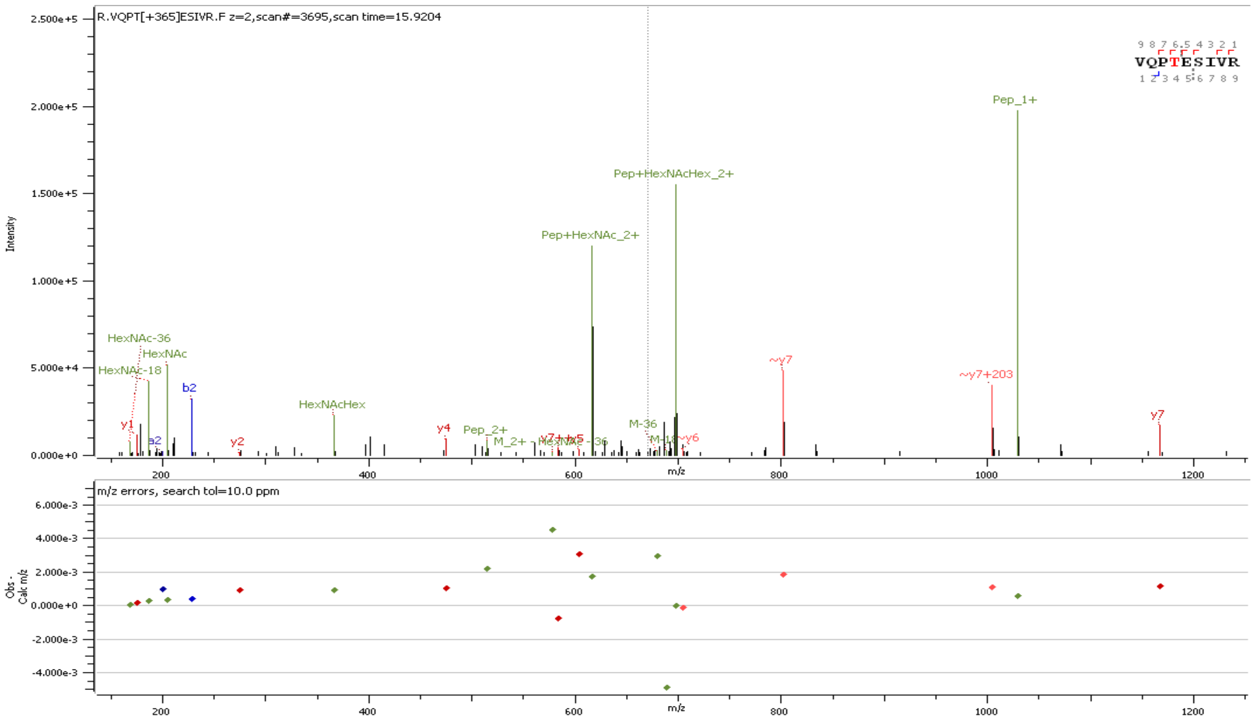


**Supp. Figure S2** Representative HCD MS/MS spectrum of intact O-glycopeptide with assigned O-glycan ($GalNAc_1Gal_1$) at T323

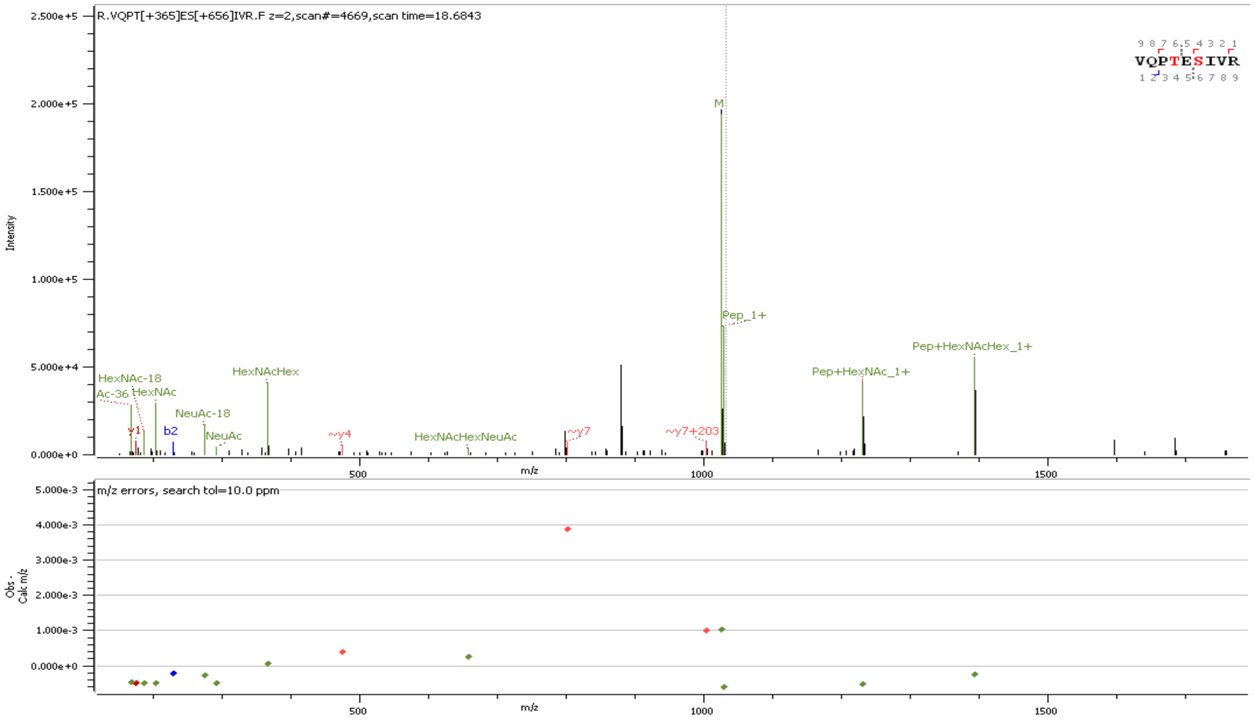

**Supp. Figure S3** Representative HCD MS/MS spectrum of intact O-glycopeptide with assigned O-glycan ($GalNAc_1Gal_1$) at T323 & ($GalNAc_1Gal_1NeuAc_1$) at S325

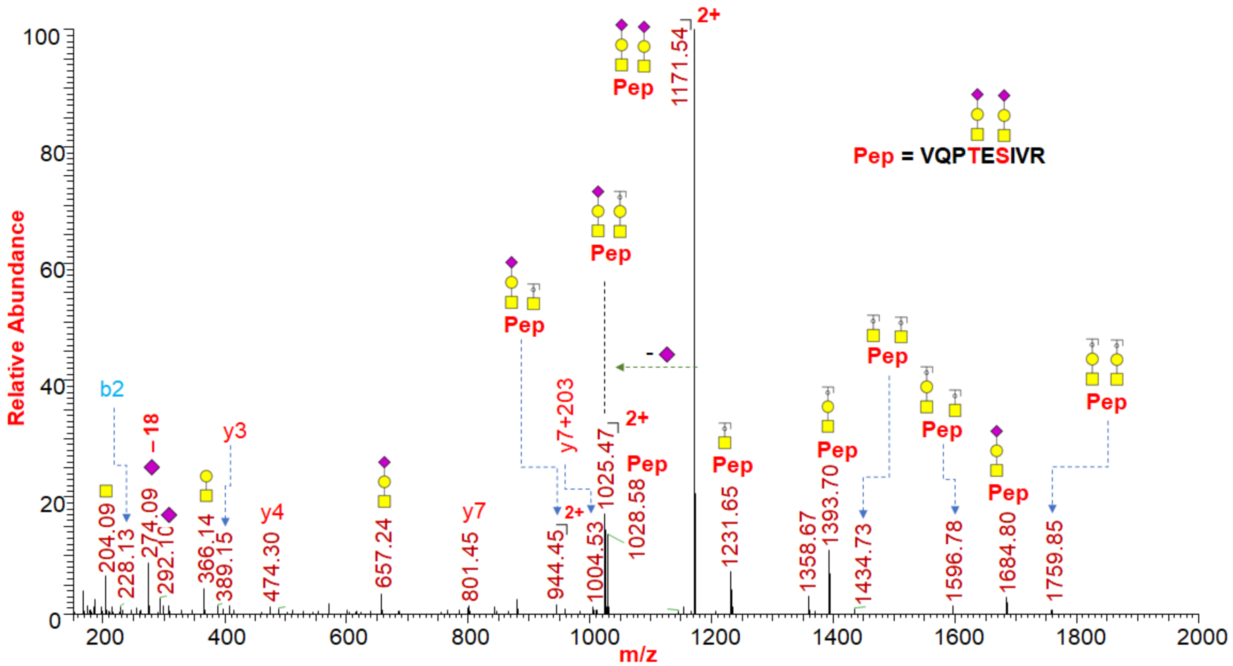

**Supp. Figure S4** Representative HCD MS/MS spectrum of intact O-glycopeptide with assigned O-glycan ($GalNAc_1Gal_1NeuAc_1$) at T323 & ($GalNAc_1Gal_1NeuAc_1$) at S325

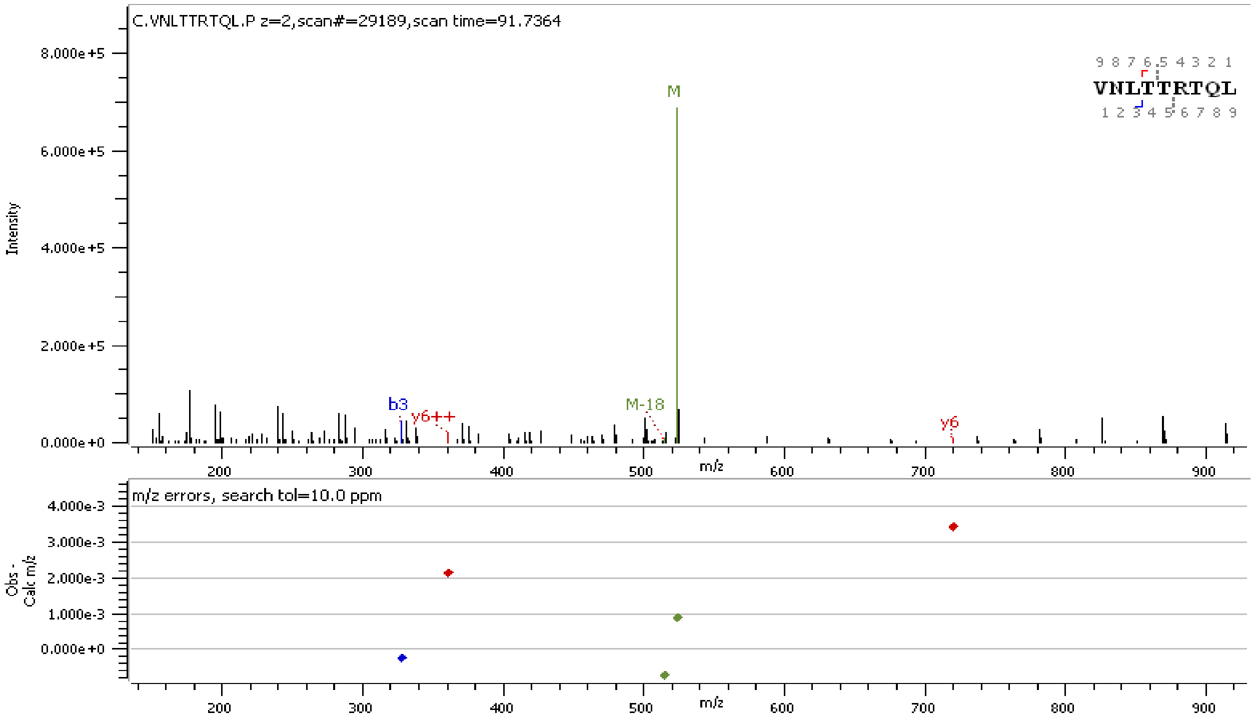


**Supp. Figure S5.** Representative HCD MS/MS spectrum of non-glycosylated peptide at N17

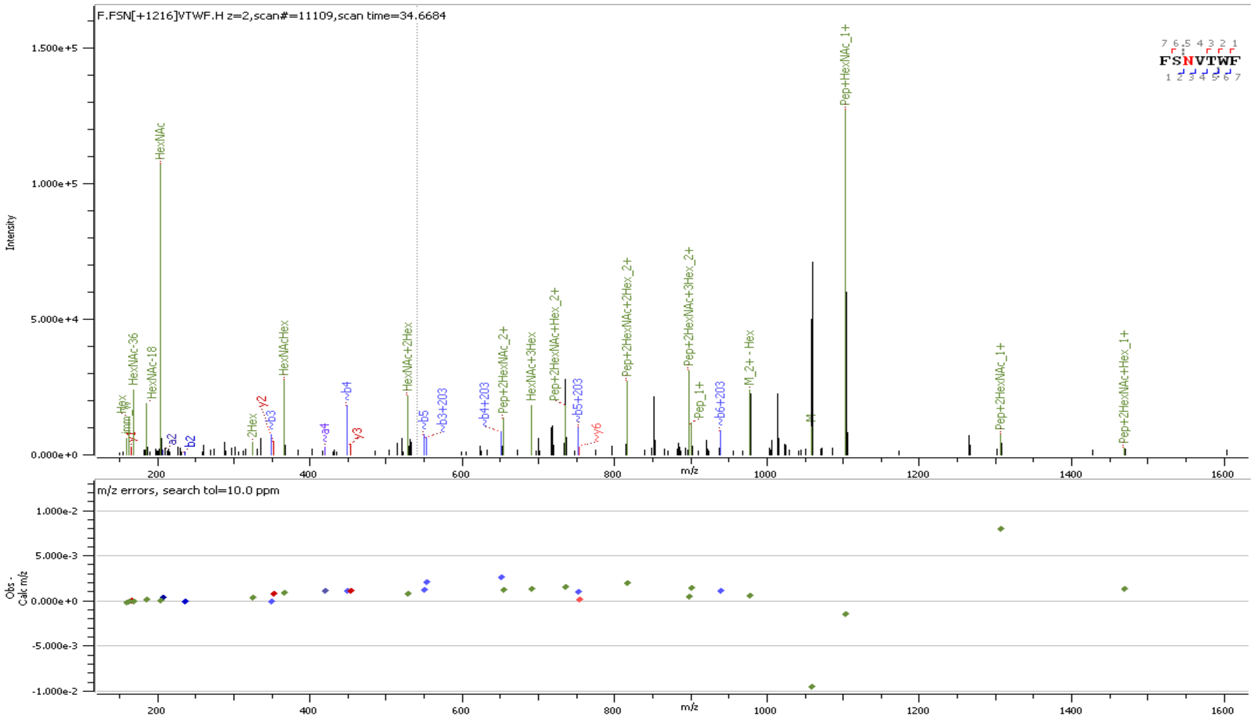


**Supp. Figure S6** Representative HCD MS/MS spectrum of intact N-glycopeptide with assigned N-glycan ($GlcNAc_2Man_5$) at N61

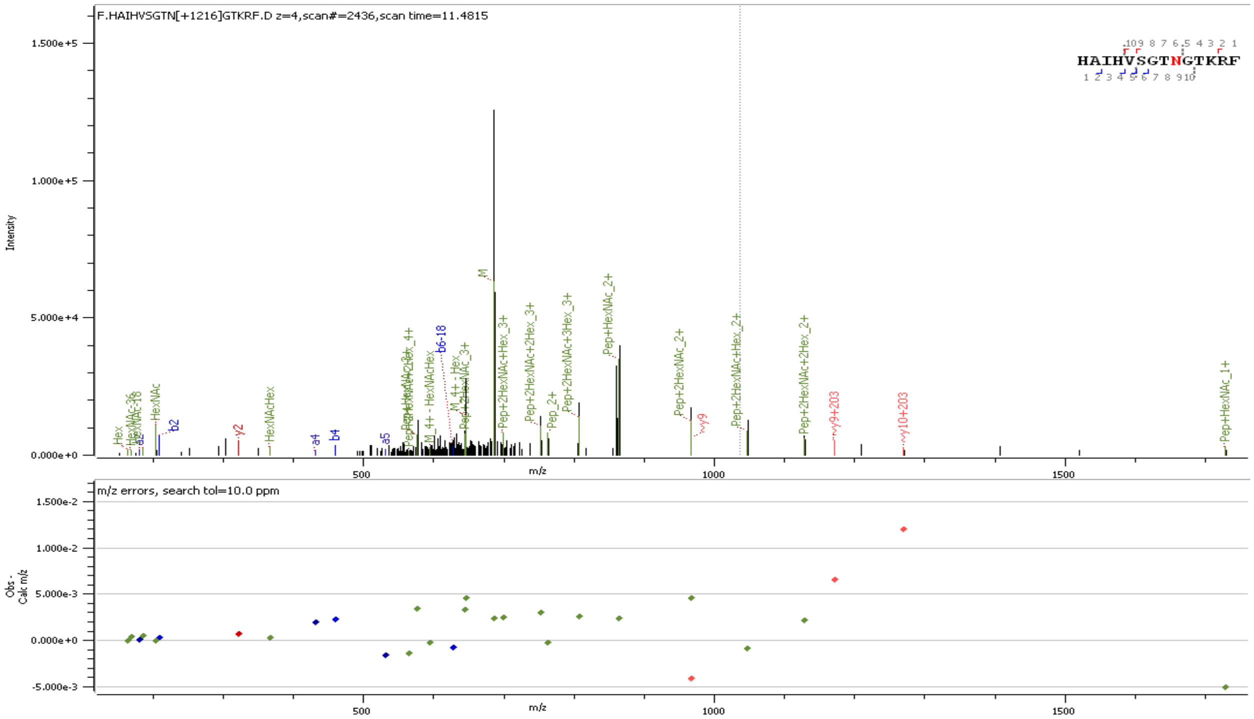


**Supp. Figure S7** Representative HCD MS/MS spectrum of intact N-glycopeptide with assigned N-glycan ($GlcNAc_2Man_5$) at N74

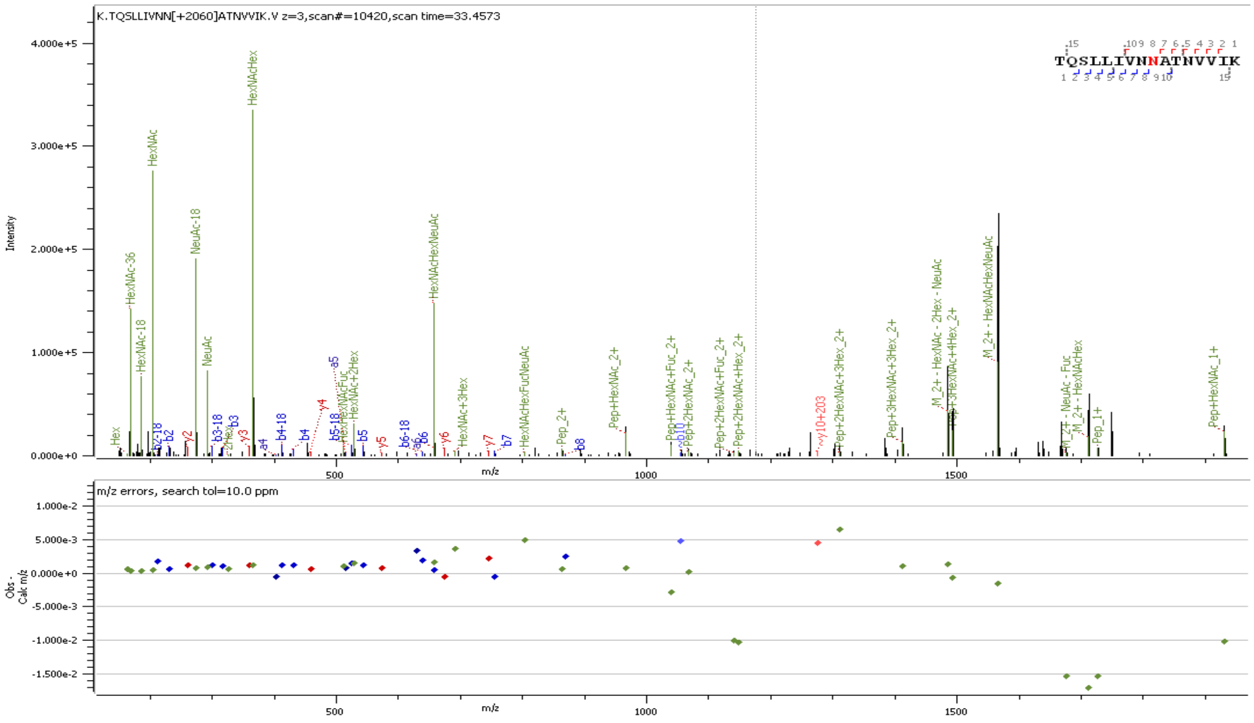


**Supp. Figure S8.** Representative HCD MS/MS spectrum of intact N-glycopeptide with assigned N-glycan ($GlcNAc_2Fuc_1Man_3GlcNAc_2Gal_1NeuAc_1$) at N122

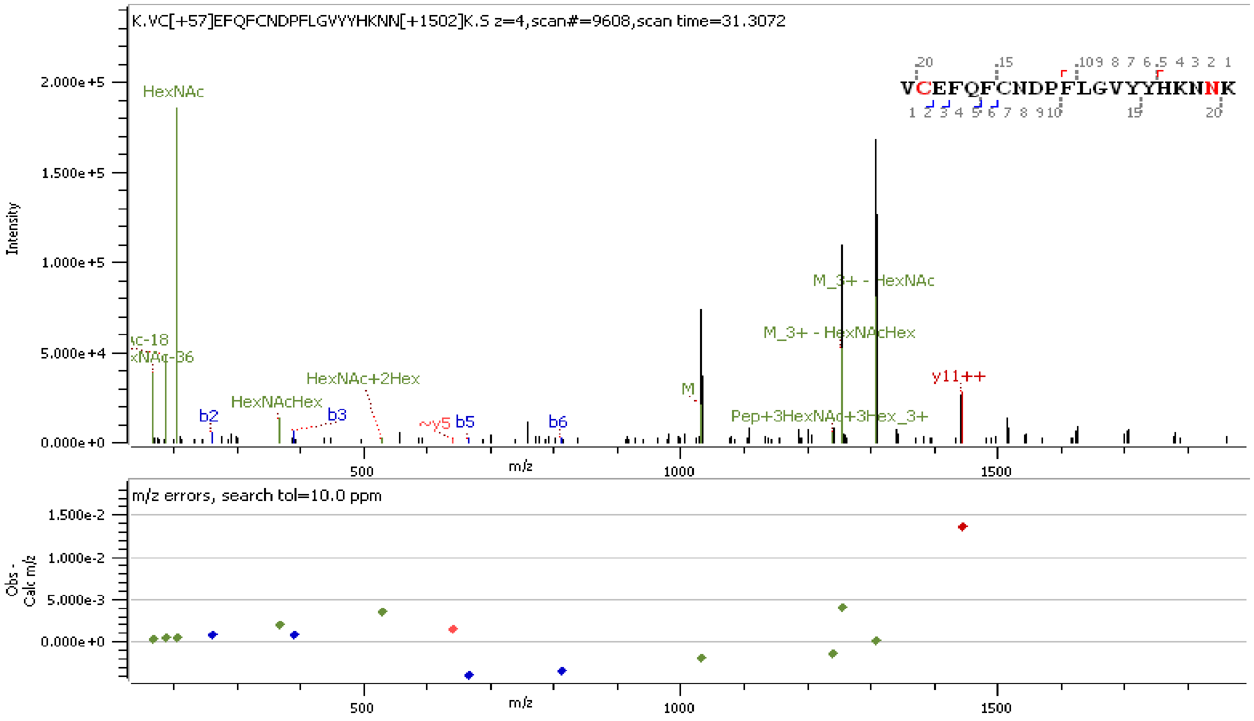


**Supp. Figure S9.** Representative HCD MS/MS spectrum of intact N-glycopeptide with assigned N-glycan ($GlcNAc_2Man_3GlcNAc_4$) at N149

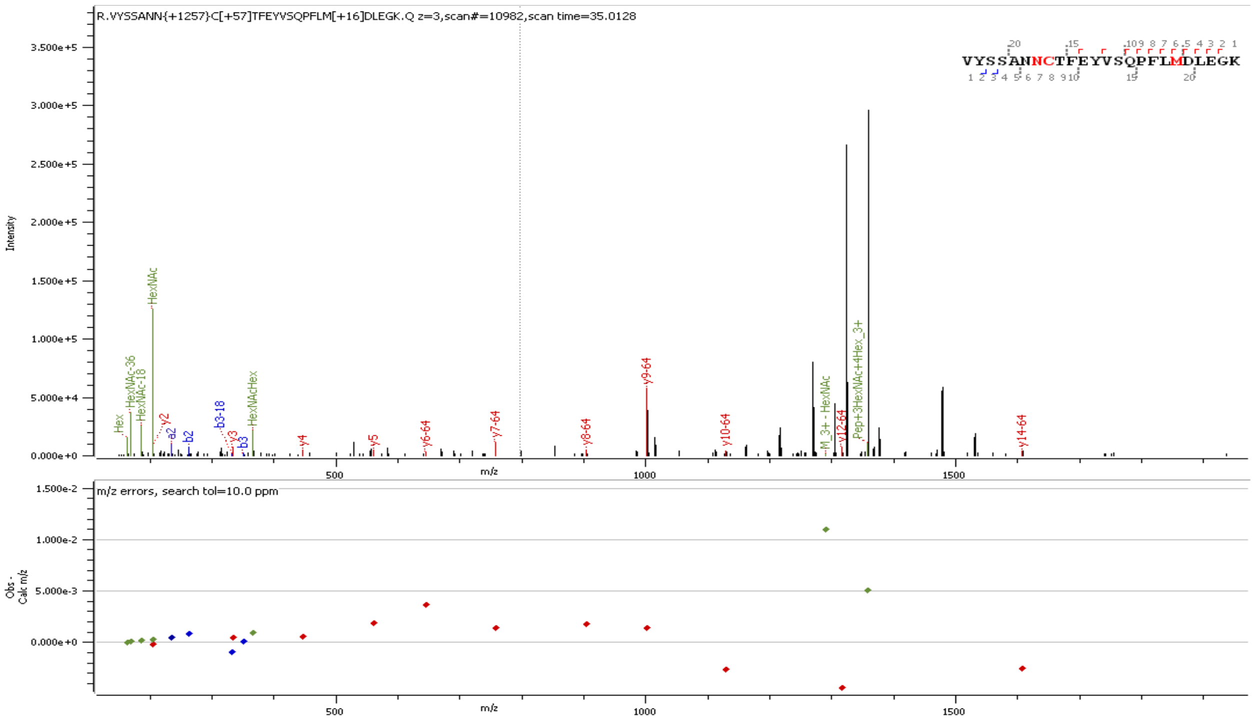


**Supp. Figure S10.** Representative HCD MS/MS spectrum of intact N-glycopeptide with assigned N-glycan ($GlcNAc_2Man_3GlcNAc_1Gal_1$) at N165

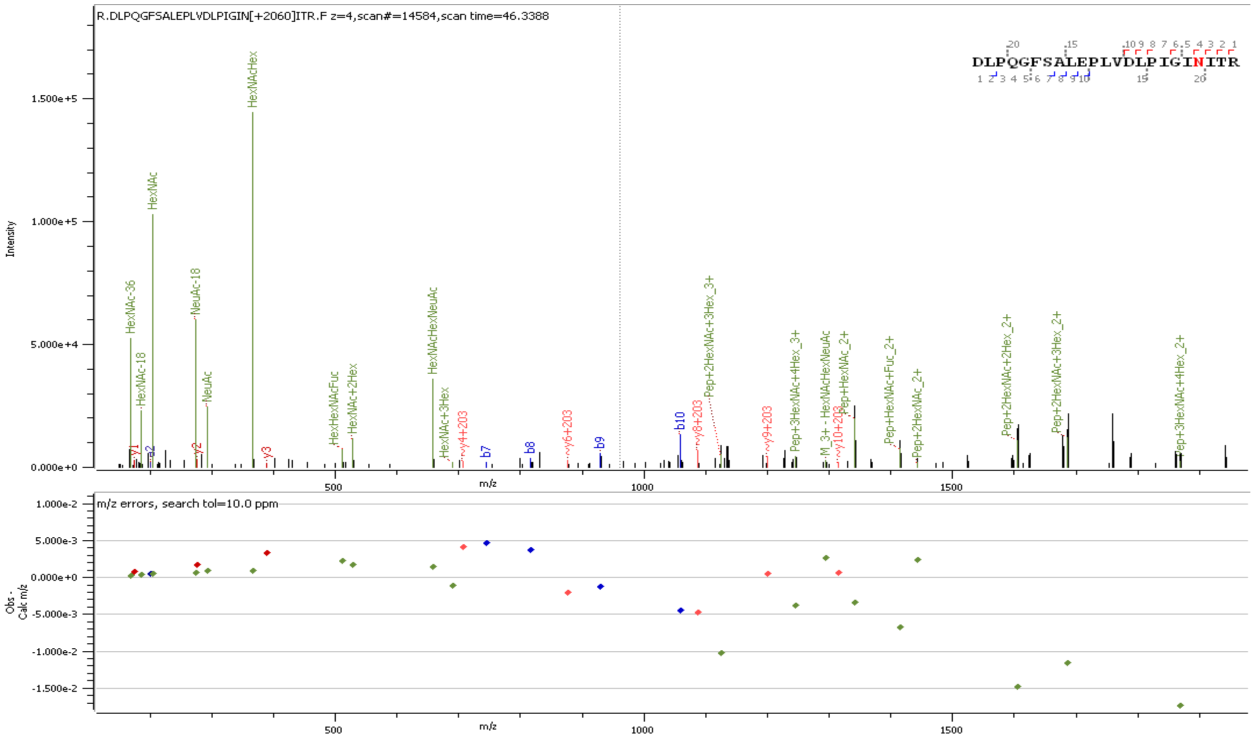


**Supp. Figure S1.** Representative HCD MS/MS spectrum of intact N-glycopeptide with assigned N-glycan ($GlcNAc_2Fuc_1Man_3GlcNAc_2Gal_2NeuAc_1$) at N234

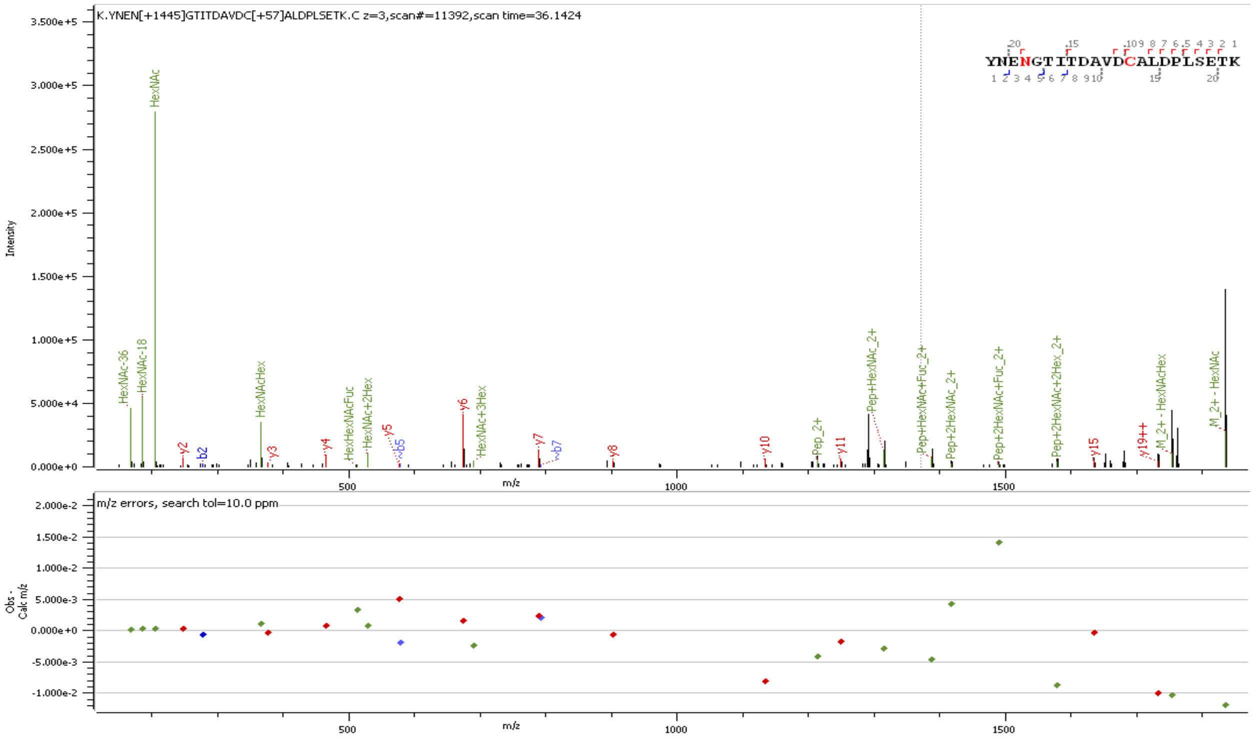


**Supp. Figure S12.** Representative HCD MS/MS spectrum of intact N-glycopeptide with assigned N-glycan ($GlcNAc_2Fuc_1Man_3GlcNAc_2$) at N282

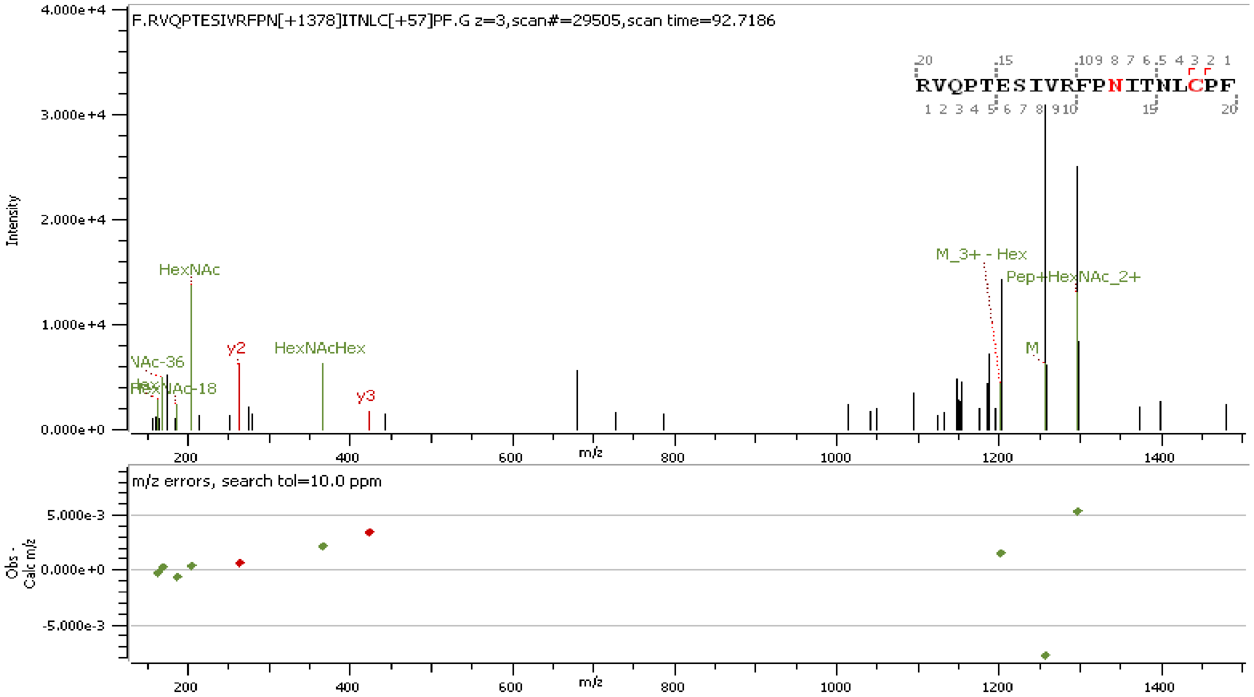


**Supp. Figure S13.** Representative HCD MS/MS spectrum of intact N-glycopeptide with assigned N-glycan ($GlcNAc_2Man_6$) at N331

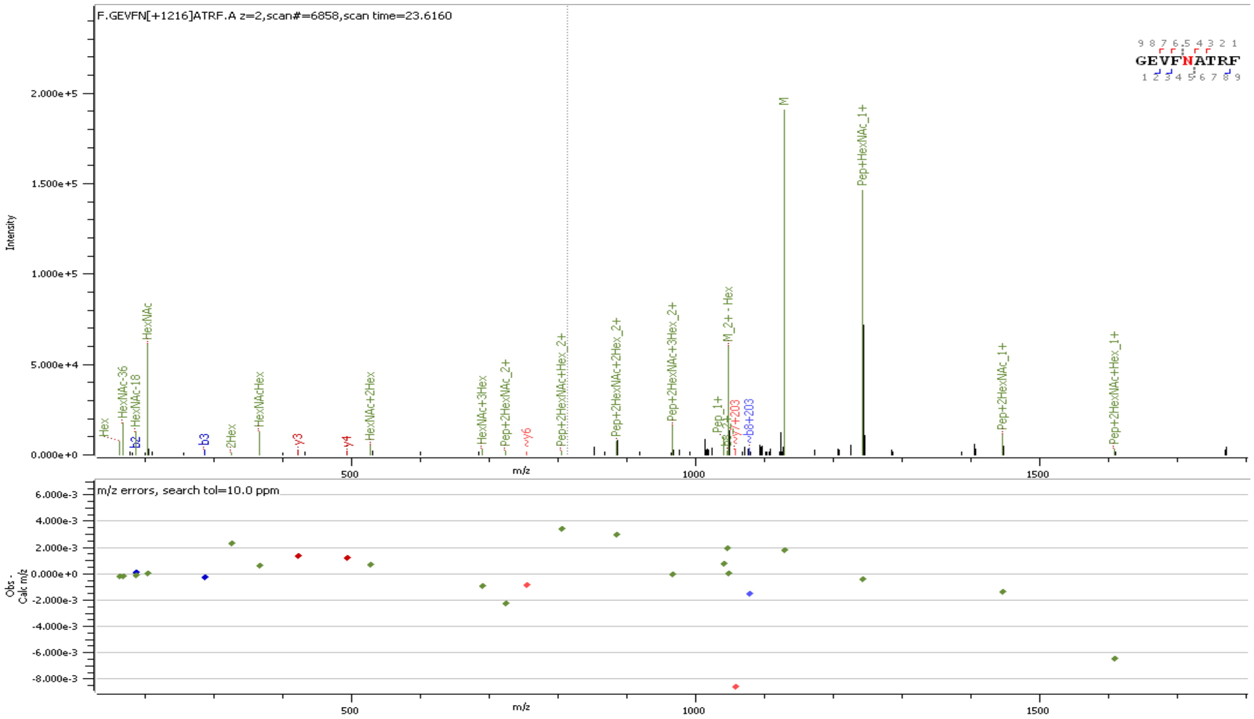


**Supp. Figure S14.** Representative HCD MS/MS spectrum of intact N-glycopeptide with assigned N-glycan ($GlcNAc_2Man_5$) at N343

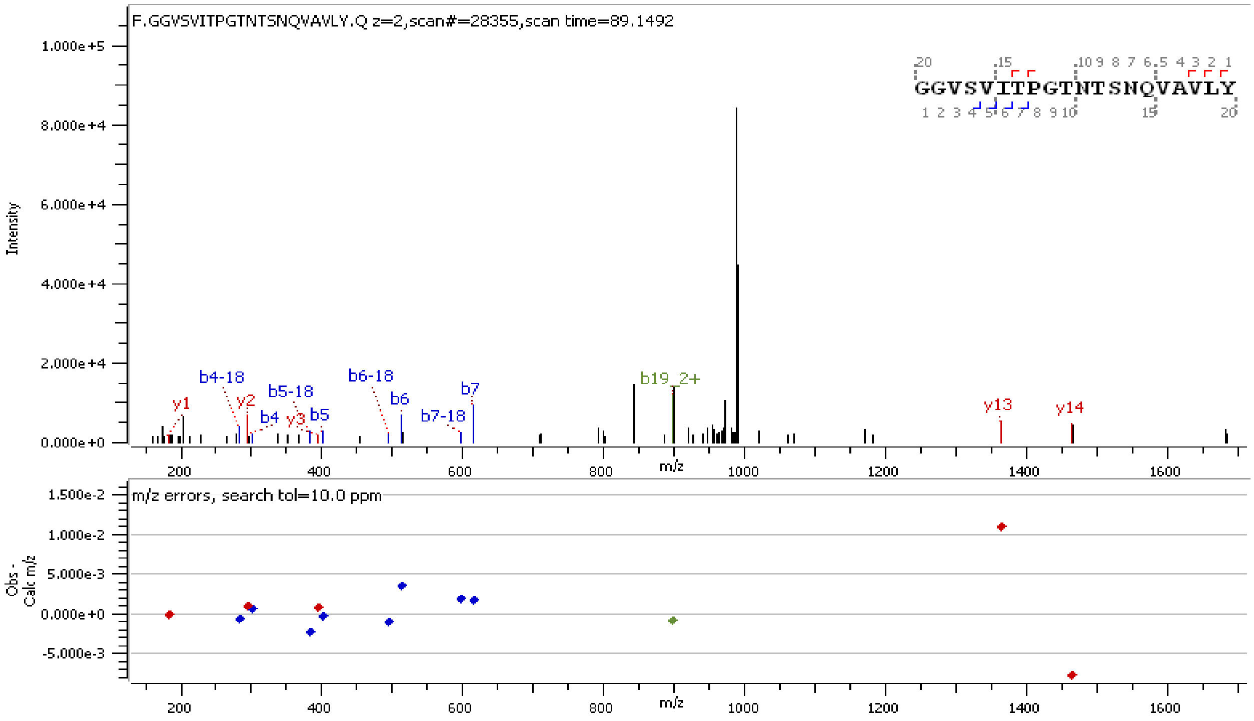


**Supp. Figure S15.** Representative HCD MS/MS spectrum of Non-glycosylated peptide at N603

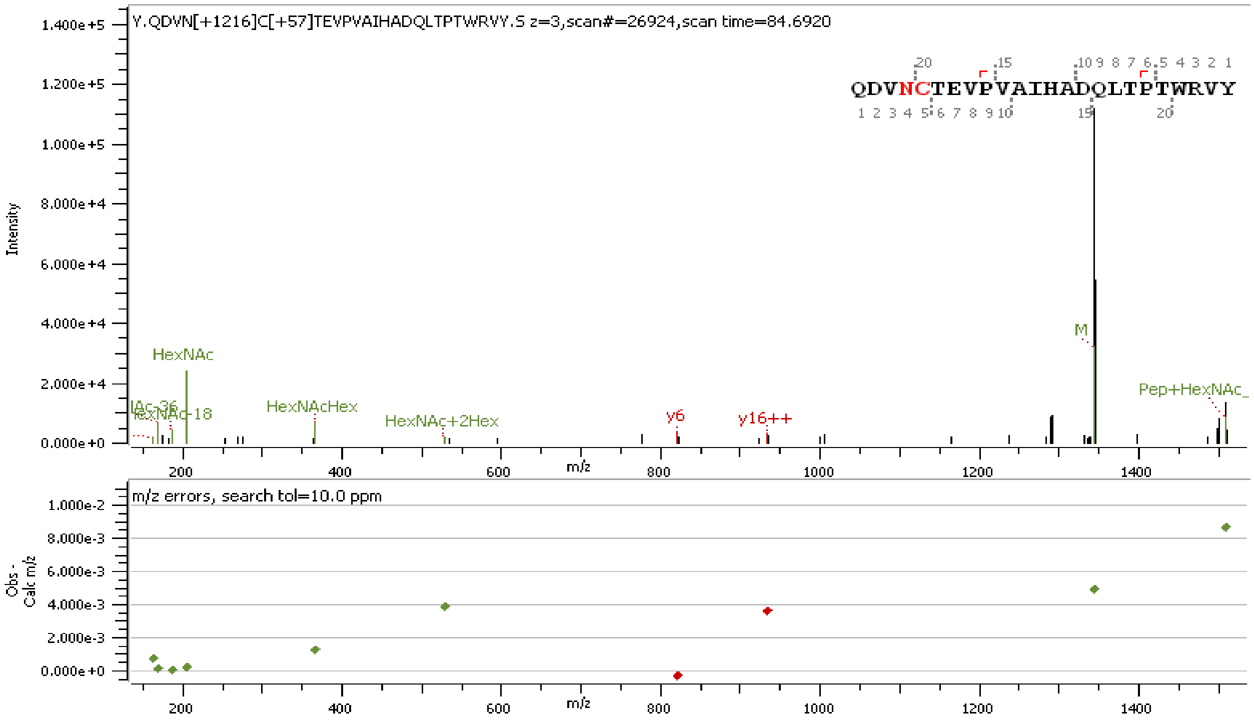


**Supp. Figure S16.** Representative HCD MS/MS spectrum of intact N-glycopeptide with assigned N-glycan ($GlcNAc_2Man_5$) at N616

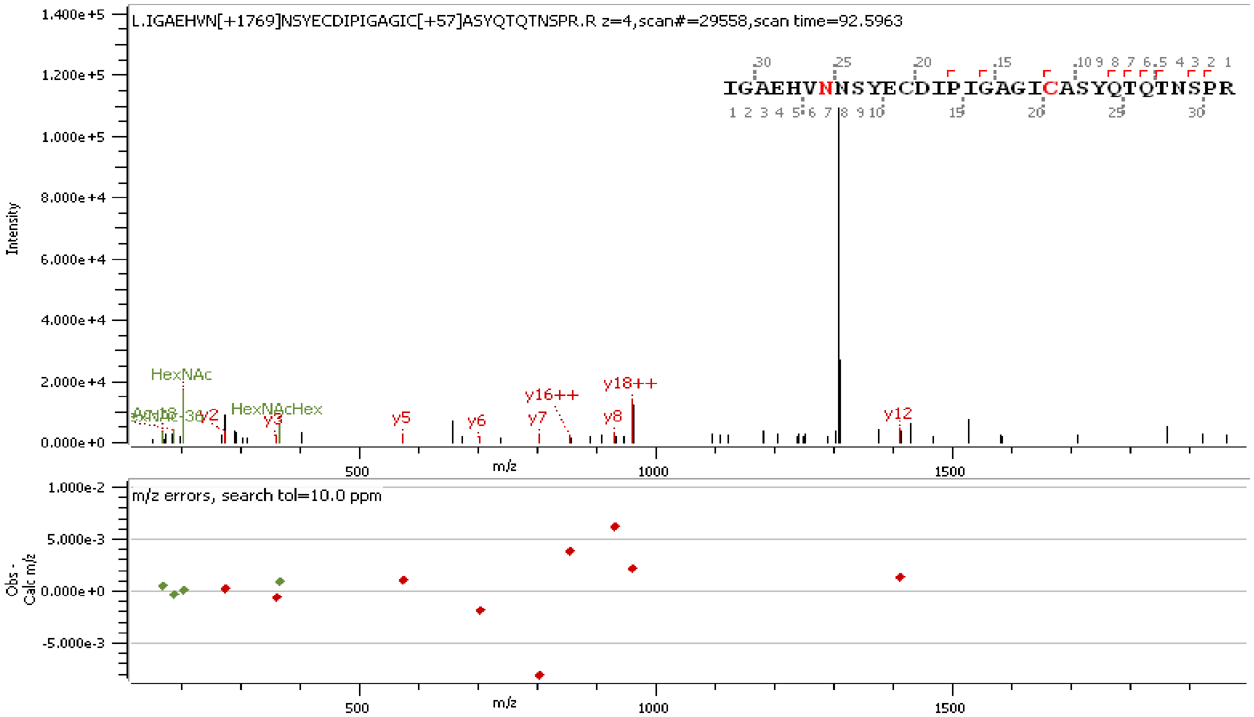


**Supp. Figure S17.** Representative HCD MS/MS spectrum of intact N-glycopeptide with assigned N-glycan ($GlcNAc_2Fuc_1Man_3GlcNAc_2$ $Gal_2$) at N657

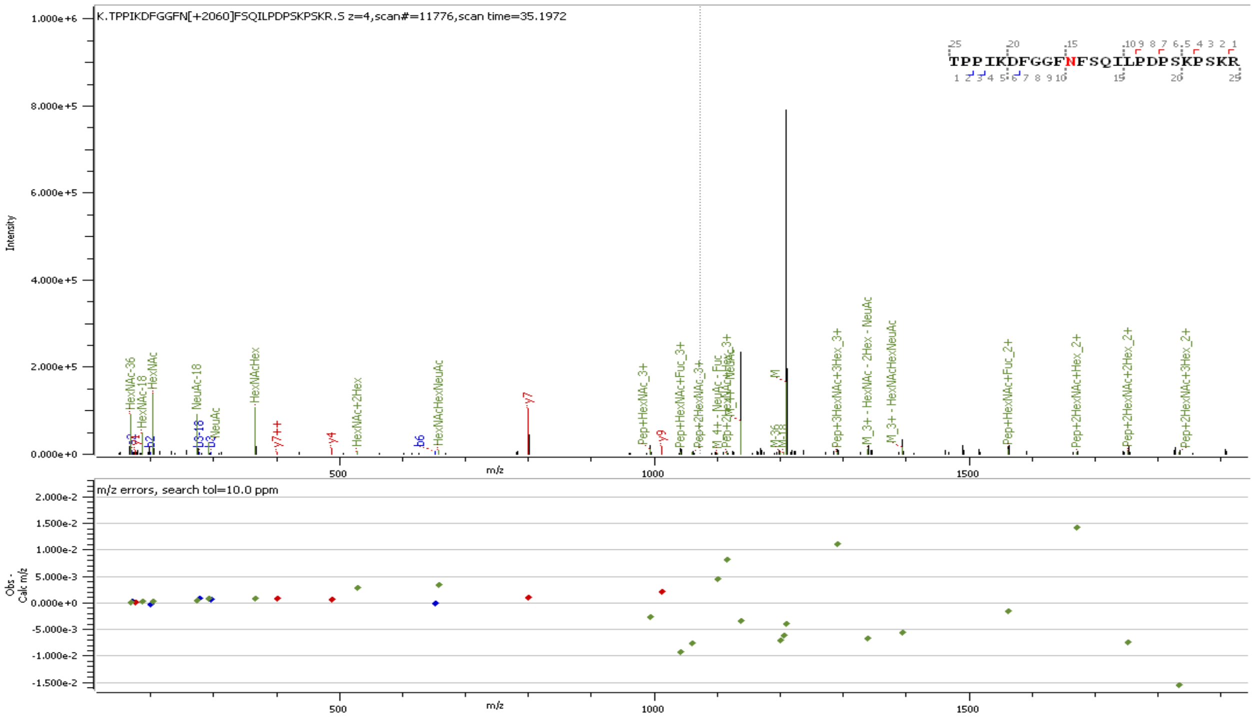


**Supp. Figure S18.** Representative HCD MS/MS spectrum of intact N-glycopeptide with assigned N-glycan ($GlcNAc_2Fuc_1Man_3GlcNAc_2Gal_2NeuAc_1$) at N801

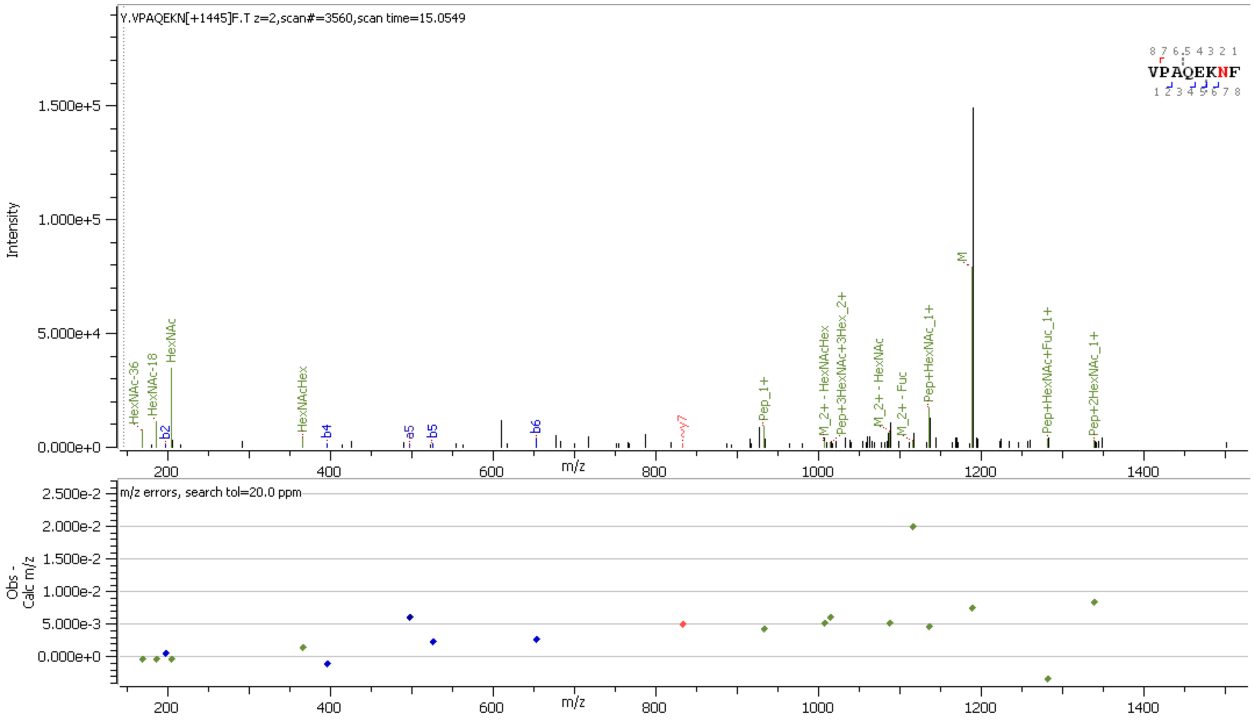


**Supp. Figure S19.** Representative HCD MS/MS spectrum of intact N-glycopeptide with assigned N-glycan ($GlcNAc_2Fuc_1Man_3GlcNAc_2$) at N1074 (peptide found to be 99.3% unglycosylated)

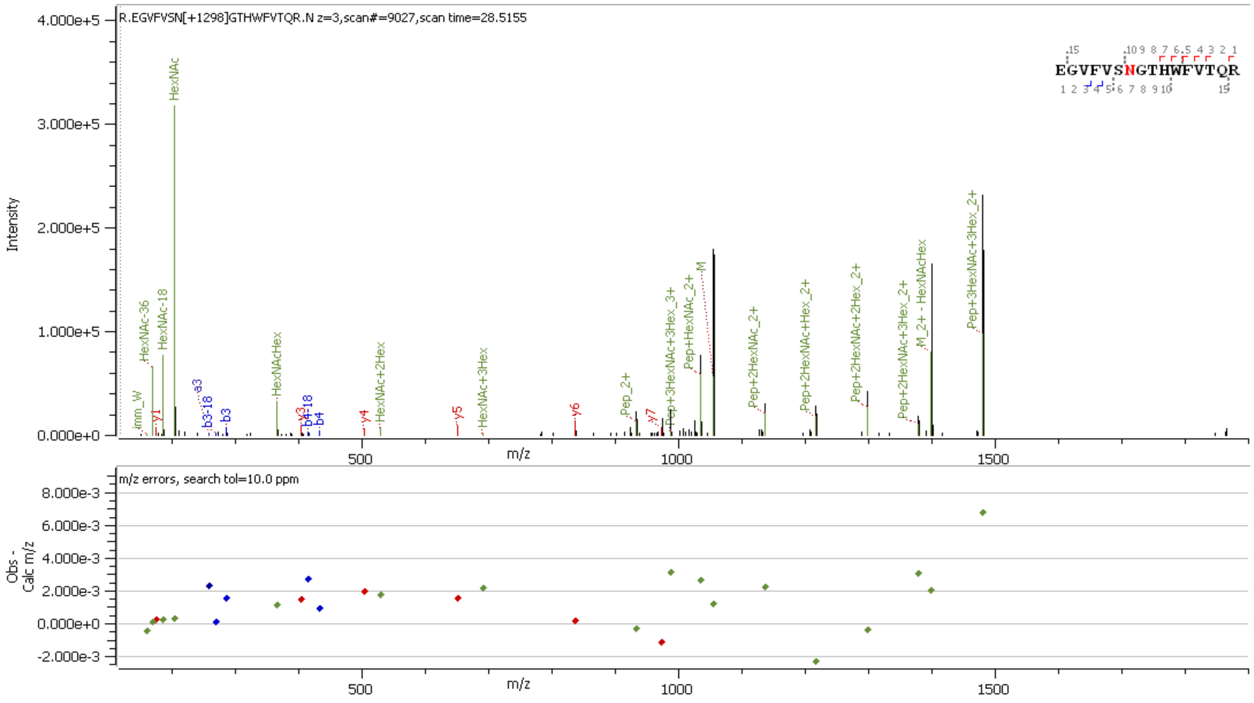


**Supp. Figure S20.** Representative HCD MS/MS spectrum of intact N-glycopeptide with assigned N-glycan ($GlcNAc_2Fuc_1Man_3GlcNAc_2Gal_2NeuAc_1$) at N1098

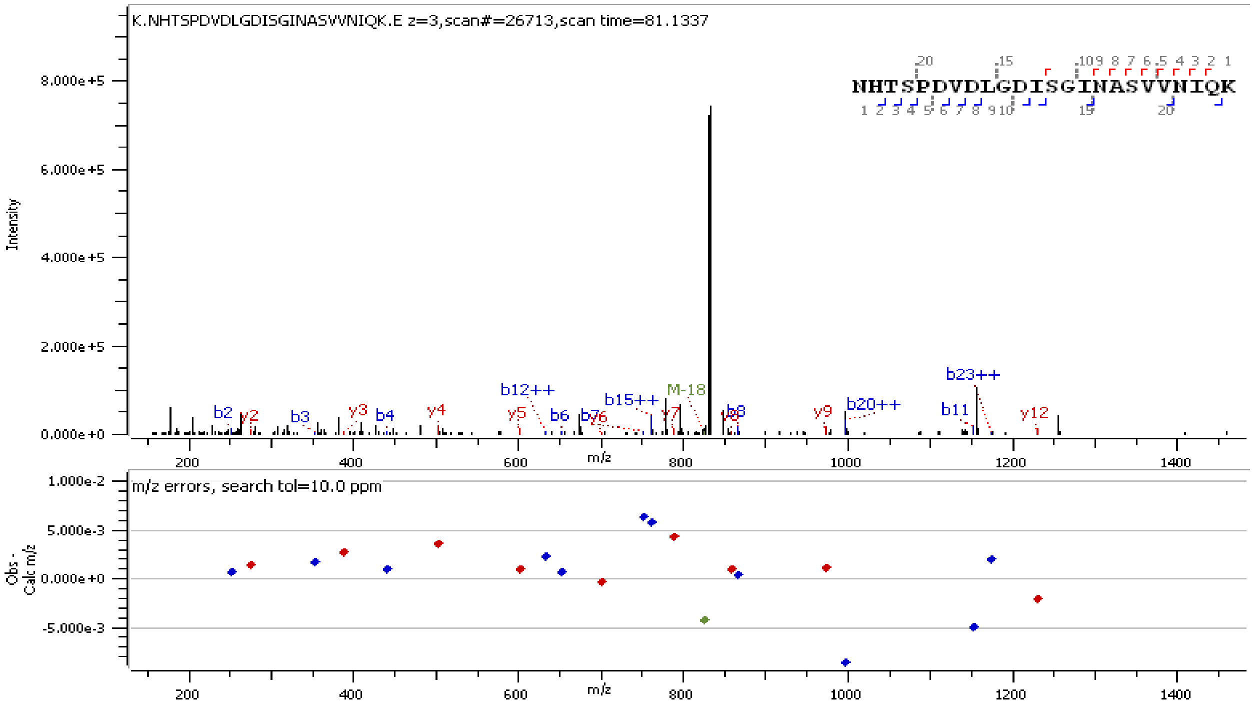


**Supp. Figure S21.** Representative HCD MS/MS spectrum of non-glycosylated peptide at N1158

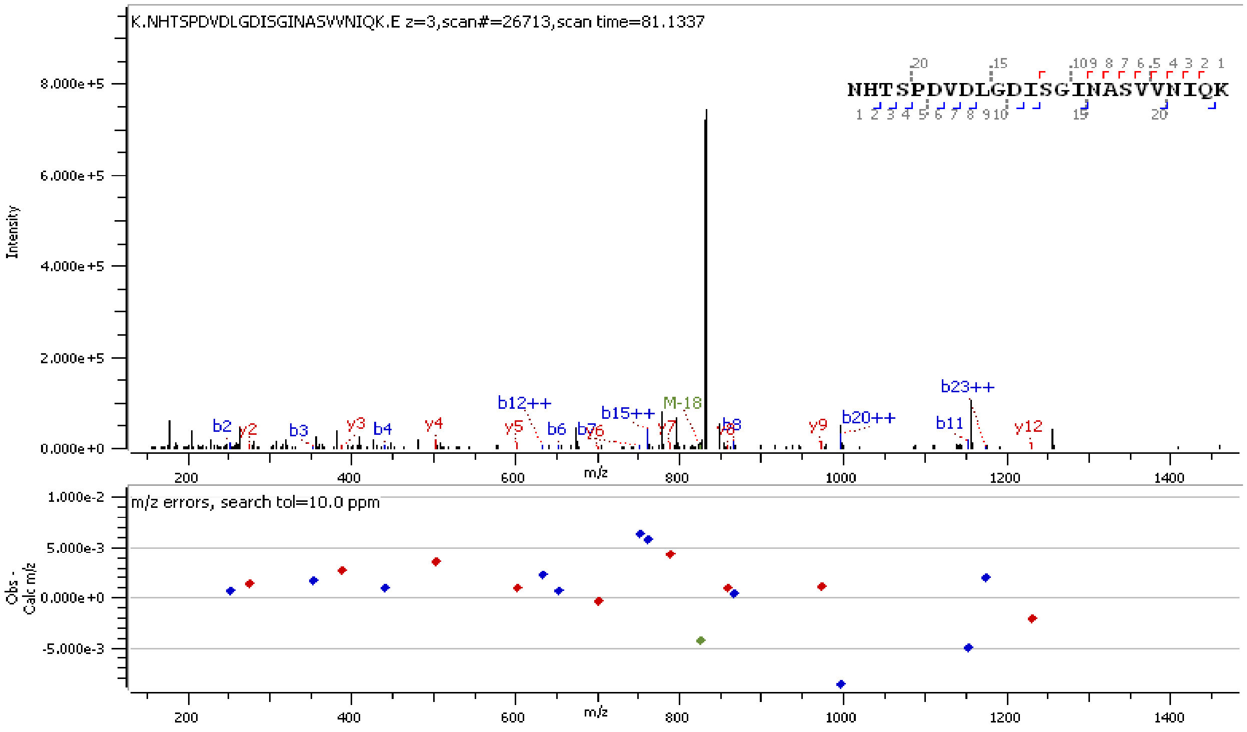


**Supp. Figure S22.** Representative HCD MS/MS spectrum of non-glycosylated peptide at N1173

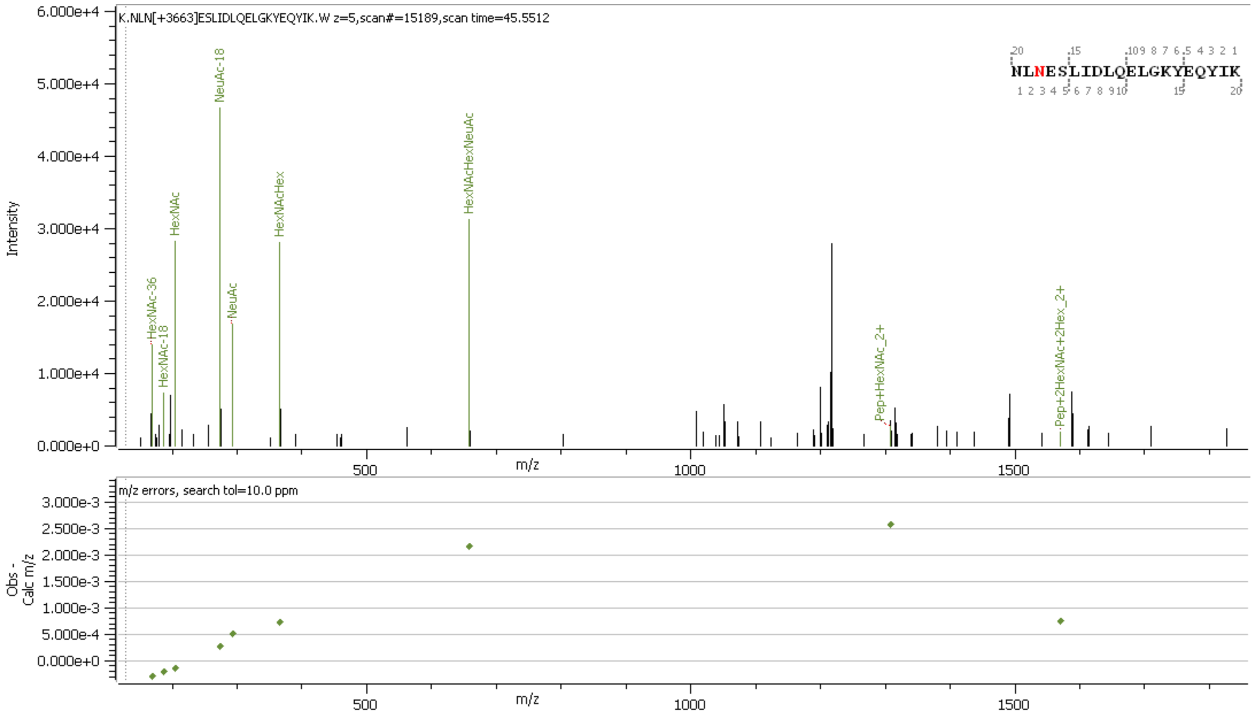


**Supp. Figure S23.** Representative HCD MS/MS spectrum of intact N-glycopeptide with assigned N-glycan ($GlcNAc_2FucMan_3GlcNAc_4Gal_4NeuAc_4$) at N1194
